# A one-step protocol to generate impermeable fluorescent HaloTag substrates for *in situ* live cell application and super-resolution imaging

**DOI:** 10.1101/2024.09.20.614087

**Authors:** Kilian Roßmann, Siqi Sun, Christina Holmboe Olesen, Maria Kowald, Eleni Tapp, Ulrich Pabst, Marie Bieck, Ramona Birke, Brenda C. Shields, PyeongHwa Jeong, Jiyong Hong, Michael R. Tadross, Joshua Levitz, Martin Lehmann, Noa Lipstein, Johannes Broichhagen

## Abstract

Communication between cells is largely orchestrated by proteins on the cell surface, which allow information transfer across the cell membrane. Super-resolution and single-molecule visualization of these proteins can be achieved by genetically grafting HTP (HaloTag Protein) into the protein of interest followed by brief incubation of cells with a dye-HTL (dye-linked HaloTag Ligand). This approach allows for use of cutting-edge fluorophores optimized for specific optical techniques or a cell-impermeable dye-HTL to selectively label surface proteins without labeling intracellular copies. However, these two goals often conflict, as many high-performing dyes exhibit membrane permeability. Traditional methods to eliminate cell permeability face synthetic bottlenecks and risk altering photophysical properties. Here we report that dye-HTL reagents can be made cell-impermeable by inserting a charged sulfonate directly into the HTL, leaving the dye moiety unperturbed. This simple, one-step method requires no purification and is compatible with both the original HTL and second-generation HTL.2, the latter offering accelerated labeling. We validate such compounds, termed dye-SHTL (‘dye shuttle’) conjugates, in live cells via widefield microscopy, demonstrating exclusive membrane staining of extracellular HTP fusion proteins. In transduced primary hippocampal neurons, we label mGluR2, a neuromodulatory G protein-coupled receptor (GPCR), with dyes optimized for stimulated emission by depletion (STED) super-resolution microscopy, allowing unprecedented accuracy in distinguishing surface and receptors from those in internal compartments of the presynaptic terminal, important in neural communication. This approach offers broad utility for surface-specific protein labelling.

## INTRODUCTION

Self-labelling protein tags, e.g. SNAP and HTP, are at the forefront of approaches to label proteins of interest bio-orthogonally with chemical moieties in living cells and tissue.^1–3^ The main application is for fluorescence microscopy, where dyes are covalently attached to HTP fusion proteins to investigate protein localization and dynamics.^4,5^ A particularly important superfamily of signalling proteins are cell surface receptors, which mediate information transmission between cells. However, standard protocols label both the extracellular and intracellular populations, including proteins in endo/lysosomal compartments and translated proteins that remain in the endoplasmic reticulum (**Fig. 1A**, left). Selective labelling of the cell-surface subset can be achieved with cell-impermeable dyes. Typically, this requires that the dye itself bears anionic residues, e.g. carboxylates and sulfonates (**Fig. 1A**, middle)^6,7^. However, many of the best dyes for single molecule and super-resolution microscopy^8–11^ are cell permeable, limiting their utility for selective surface labelling.

**Figure 1:**
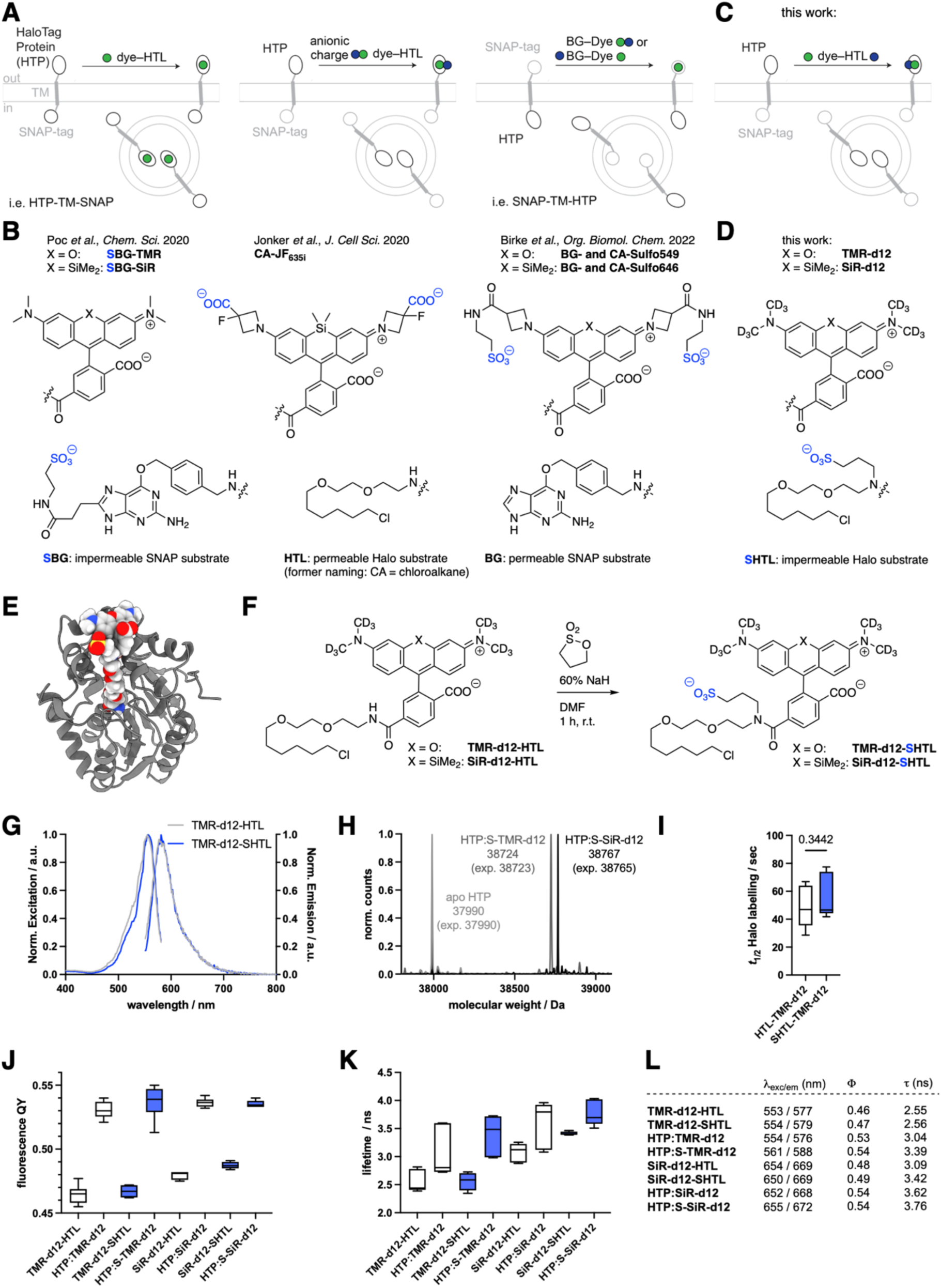
Logic for impermeable dyes to label cell surface proteins. **A-B)** Known strategies to address extracellularly exposed self-labelling tags with cartoons and chemical structures indicating where anionic charges have previously been introduced. **C-D)** Our new strategy to install a sulfonate charge on the HaloTag Ligand (HTL). **E)** Modelling of the HaloTag Protein (HTP) bound to TMR-SHTL. **F)** Synthesis of TMR-d12-SHTL and SiR-d12-SHTL. **G)** Excitation and emission profiles of TMR-d12 comparing HTL to SHTL conjugates. **H)** *In vitro* protein labelling of apo-HTP confirms binding by full protein mass spectrometry. **I)** Labelling kinetics of apo-HTP with TMR-d12-HTL and TMR-d12-SHTL. **J)** Fluorescence quantum yields of TMR/SiR-d12 conjugated to HTL/SHTL with or without HTP-bound. **K)** Fluorescence lifetimes of the same reagents. **L)** Table summarizing values from G, J, K.

Traditionally, dyes are made less cell permeable by custom modification to the dye itself. For instance, Jonker et al., added carboxylates on the 3-position of the azetidine of JaneliaFluor635 (JF_635_), yielding impermeable and fluorogenic JF_635i_ for investigating endocytotic turnover^6^ (**Fig. 1A, B**), and Eiring et al. reported benefits for single-molecule localization microscopy with a highly charged Cy5b^10^. Our own laboratories have addressed this in a similar vein, by fusing a sulfonate on the same azetidine position via amide bond coupling using taurine, effectively converting JF_549_ and JF_646_ to Sulfo549 and Sulfo646 (ref ^7^), respectively (**Fig. 1A**, right and **Fig. 1B**), which have been used to study kainate receptor stoichiometry^12^ and intracellular trafficking properties of GPCR subtypes^13^.

A more general and effective strategy would offer a universal solution for all dyes. For instance, we recently described a modified version of the SNAP ligand in which the benzylguanine (BG) leaving group carries a negative charge due to incorporation of a sulfonate on the 8-position of guanine, resulting in ‘SBG substrates’ that are released upon covalent reaction (**Fig. 1A**, right and **Fig. 1B**).^14^ Unfortunately, this approach is not possible for the HaloTag system, since the HTL leaving group is a chloride anion that cannot be chemically modified.

In this study, we develop a simple approach for late-stage introduction of a sulfonate on the amide bond that links HTL to the dye (**Fig. 1C**). We determine the photophysical properties of the subsequent modified dyes, validate their lack of membrane permeability in cell lines and primary neurons, and devise a protocol to quantitatively convert commercially available JaneliaFluor-HTL reagents—and potentially other dyes—to enable straightforward, widespread application.

## RESULTS

In this work, we aimed to introduce a sulfonate on the amide bond of available dye-HTL conjugates (**Fig. 1C, D**), giving access to dye-SHTL substrates. We first used molecular modelling to assess if this would be tolerated by the HTP, using AutoDock 4 engine alongside the two-point attraction method for covalent docking^15,16^ on the TMR-HTL bound HTP structure (PDB-ID 6Y7A)^17^. We found that the added C3-linker bearing the sulfonate does not sterically hamper the exposed dye on the protein surface (**Fig. 1E**) (see SI and **Fig. S1-S4**). Next, we synthesized two dye-SHTL substrates starting with our recently reported TMR-d12 and SiR-d12 dyes, the latter of which has been shown to be an outstanding candidate for STED super-resolution microscopy.^18^ Dissolving these dye-HTL substrates in DMF and subsequently adding sodium hydride (NaH 60% in mineral oil) before 1,3 propane sultone, led to clean conversion, yielding TMR-d12-SHTL and SiR-d12-SHTL within an hour in 94% and 91% yield, respectively, after HPLC purification (**Fig. 1F**) (see SI). Introduction of the charged sulfonate only slightly changed the excitation and emission profiles, as illustrated for TMR-d12 (HTL: α_Ex/Em_ = 553/577 nm; SHTL: α_Ex/Em_ = 554/579 nm) (**Fig. 1G, L**). Full protein mass spectrometry on recombinant HTP (ref ^7^) confirmed that TMR-d12-SHTL and SiR-d12-SHTL bind to HTP with quantitative stoichiometry (**Fig. 1H**), with no significant difference in labelling kinetics as determined by fluorescence polarization (*t*_1/2_ ∼ 50 sec) (**Fig. 1I**). We next measured quantum yields (**Fig. 1J, L**) and fluorescence lifetimes (**Fig. 1K, L**) to test if sulfonation impairs dye photophysics. The same trend in increasing quantum yield upon binding to HTP was observed for all dyes (Φ_HTP-free_ = 47–49%; <λ_HTP-bound_ ∼ 53%), while the sulfonated versions displayed up to 11% longer fluorescence lifetimes over non-sulfonated precursors (**Fig. 1L**).

We next tested both dyes in HEK293 cells transiently transfected with our previously reported SNAP-TM-HTP and HTP-TM-SNAP constructs^7^. Having each tag residing on the opposite side of the cellular membrane separated by a single transmembrane (TM) domain enabled us to simultaneously assess membrane permeability and labelling with built-in controls for expression and cell health. We first used HTP-TM-SNAP cells, using the cell-permeable BG-JF_646_ to label all intracellular proteins, co-applied for 30 minutes with either TMR-d12-HTL (**Fig. 2A**, upper row) or TMR-d12-SHTL (**Fig. 2A**, lower row). As expected, TMR-d12-HTL showed considerable intracellular signals (**Fig. 2B**) confirmed by the similarity of line-scan profiles to the BG-JF_646_ intracellular reference (**Fig. 2C**, raw fluorescence left and normalized right). By contrast, TMR-d12-SHTL (**Fig. 2E**) exhibited selective surface labelling, with line scans differing considerably from the intracellular reference. We then swapped the self-labelling tags with the SNAP-TM-HTP construct, switching to a cell-impermeable BG-SulfoJF_646_ so that the SNAP label serves as a surface-exclusive reference. Consistent with our prior findings, TMR-d12-HTL (**Fig. 2F**, upper row) exhibited intracellular labelling (**Fig. 2G, H**), while TMR-d12-SHTL (**Fig. 2F**, lower row) showed no detectable labelling (**Fig. 2I, J**). The same experiment was conducted with SiR-d12 fused to HTL (**Fig. 2K-O**) vs. SHTL (**Fig. 2P-T**) and respective red fluorophores (BG-JF_549_ and BG-Sulfo549), confirming that SHTL shows no membrane permeability regardless of the fluorophore. We further titrated SiR-d12 HTL and SHTL conjugates up to 5000 nM on HTP-TM-SNAP expressing HEK293 cells, and again observed clear membrane staining for sulfonated versions (**Fig. S5**)

**Figure 2:**
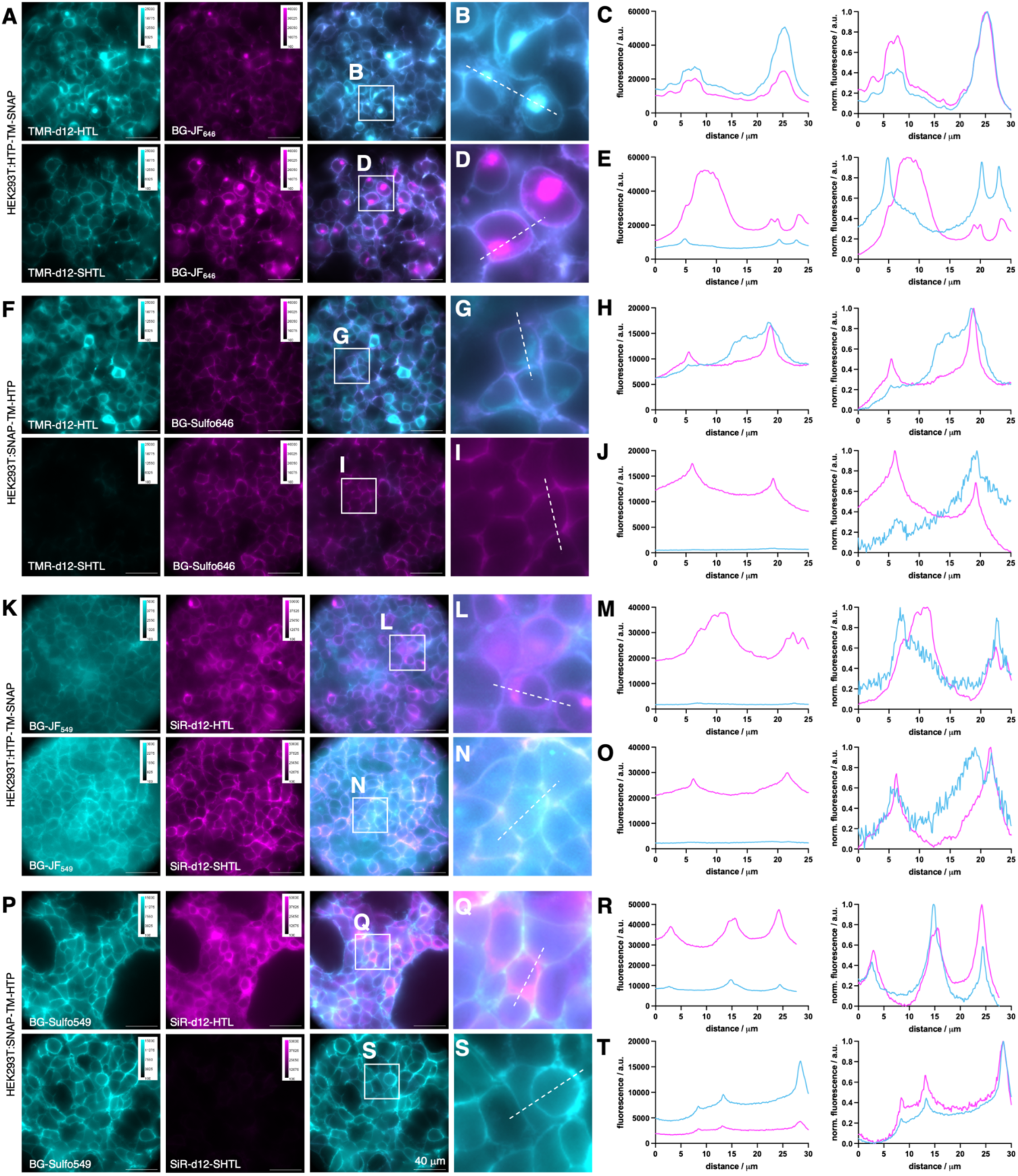
Live cell imaging in transfected HE293 cells. **A-E)** HEK293 expressing HTP-TM-SNAP. Intracellular SNAP labelled with BG-JF_646_; extracellular HTP labelled with TMR-d12-HTL or TMR-d12-SHTL. Widefield imaging (A), zoom-ins (B, D) and line scans (C, E). **F-J)** HEK293 cells expressing SNAP-TM-HTP. Extracellular SNAP labelled with BG-SulfoJF_646_; intracellular HTP labelled with TMR-d12-HTL or TMR-d12-SHTL. Widefield (F), zoom-in (G, I) and line scans (H, J). **K-O)** As for A-E but staining with BG-JF_549_ and SiR-d12-(S)HTL. **P-T)** As for F-J but staining with BG-SulfoJF_549_ and SiR-d12-(S)HTL.

We next used lentiviral particles to transduce primary mouse hippocampal neurons with a HTP-fused metabotropic glutamate receptor 2 (HTP-mGluR2) (**Fig. 3A**), which has been shown to maintain identical glutamate sensitivity^19^ and trafficking^13^ compared to untagged receptors. mGluR2 is a family C member of the G protein-coupled receptors involved in modulation of synaptic transmission and serves as a potential target for the treatment of neurological and psychiatric disorders^20^. The endogenous localization of mGluR2 is along axons and presynaptic sites^21^, although the precise sub-synaptic localization is not fully understood.^20,22^ We aimed to determine the subcellular HTP-mGluR2 distribution by antibody staining against known markers of dendritic (MAP2), presynaptic (Bassoon) and postsynaptic (Shank) sites (**Fig. 3B**). One week after transduction, we applied 500 nM of SiR-d12 (-HTL or -SHTL) to live cells for 30 minutes before fixation and imaging on a confocal microscope. The HTL variant gave rise to pronounced signals stemming from intracellular sites in the soma (**Fig. 3C**, and **Fig. S6C**), quantified by defining regions of interest around cell bodies, localized by terminating dendritic MAP2 staining, and calculating the mean intensity. Non-transduced neurons did not show SiR labelling (**Fig. S6A, B**), indicating that SiR-d12-HTL selectively labels HTP-mGluR2 in all cellular compartments. By contrast, with SiR-d12-SHTL, the fluorescent signal was significantly reduced in the soma (**Fig. 3D**), with all remaining fluorescence appearing to stem from the cell surface, and in processes that were either MAP2 positive (**Fig. 3E**) or negative (**Fig. 3F**), revealing that a substantial intracellular population of HTP-mGluR2 likely exists in all 3 sites.

**Figure 3:**
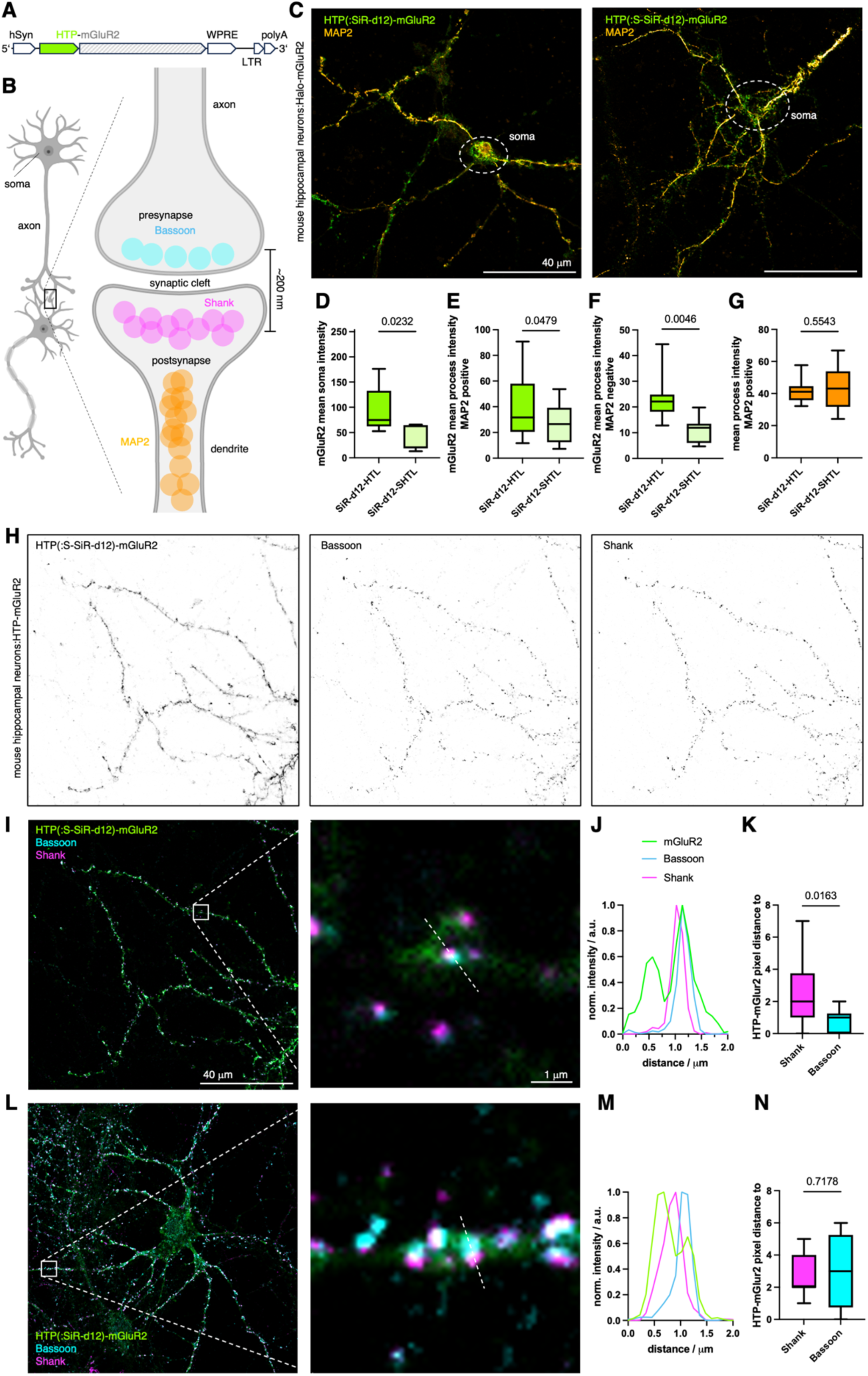
Revealing HTP-mGluR2 localization in hippocampal neurons. **A)** Viral DNA expression cassette with HTP-mGluR under hSyn promoter. **B)** Neural connection via synapses and localization of axonal MAP2, presynaptic Bassoon and postsynaptic Shank proteins (created in biorender.com). **C)** Confocal imaging of HTP-mGluR2 transduced mouse hippocampal neurons with SiR-d12-SHTL (left) and SiR-d12-HTL (right), co-stained with an antibody against MAP2 for dendrite identification. **D-G)** Quantification of HTP-mGluR2 labelling in the soma (D), in dendrites (MAP2 positive, E) and in axons (MAP2 negative, F) reveals significantly less signal using SiR-d12-SHTL, while no difference in axonal MAP2 intensity is observed (G). Mean±SD. Student’s t-test. **H)** Confocal imaging of HTP-mGluR2 transduced mouse hippocampal neurons cells with SiR-d12-SHTL (500 nM), and the pre- and postsynaptic markers Bassoon and Shank, respectively. **I)** Overlay of images in H. **J)** Representative line scan of a synapse shows mGluR2 co-localization primarily with the presynaptic marker Bassoon. **K**) Quantification of mGluR2 localization with respect to Bassoon and Shank. Mean±SD. Student’s t-test. **L-M**) As for I-K but labelling with SiR-d12-HTL.

As an important control, anti-MAP2 intensities were identical in SiR-d12-HTL and SiR-d12-SHTL samples (**Fig. 3G**). Of note, MAP2 is known to locate in dendrites and not in axons, and most synapses in this preparation are axo-dendritic. To identify synapses, antibodies against Bassoon (presynaptic marker) and Shank (postsynaptic marker) were applied (**Fig. 3B, H**) alongside SiR-d12-SHTL. Merging the channels (**Fig. 3I**) to perform line scans revealed a tendency towards pre-synaptic localization of HTP-mGluR2 with Bassoon (**Fig. 3J**), as revealed by plotting pixel distances of the respective intensity maxima (**Fig. 3K**). Such experiments outperform SiR-d12-HTL treated preparations (**Fig. 3L**), where signals likely include receptors in intracellular sites (**Fig. 3M**), therefore rendering synaptic allocation less precise, i.e. non-significant when comparing pixel distance (**Fig. 3N**, and **Fig. S6D, E**). This underlines the critical impact of true surface labelling to interrogate glutamate-responsive protein pools, since distances to release sites have been shown to be important for neural activity^23^.

After using confocal microscopy, we next turned to stimulated emission by depletion (STED) super-resolution imaging on these samples (**Fig. 4A**), keeping in mind that SiR-d12 is an excellent dye for this imaging technique^18^. Probing processes by a line scan revealed the extracellular HTP-mGluRs with a resolution of 134 nm (**Fig. 4B**) with full width at half maxima (FWHM) values of 60 and 68 nm (cf. confocal: FWHM = 259 nm). Furthermore, having a red fluorophore linked antibody targeted by immunohistochemistry against Bassoon, we were able to perform two-color STED on the contact site of a presynaptic site onto the process (**Fig. 4C**). This demonstrated the usefulness of addressing distinct pools of proteins in combination with super-resolution imaging, gaining information of neural ultrastructures.

**Figure 4:**
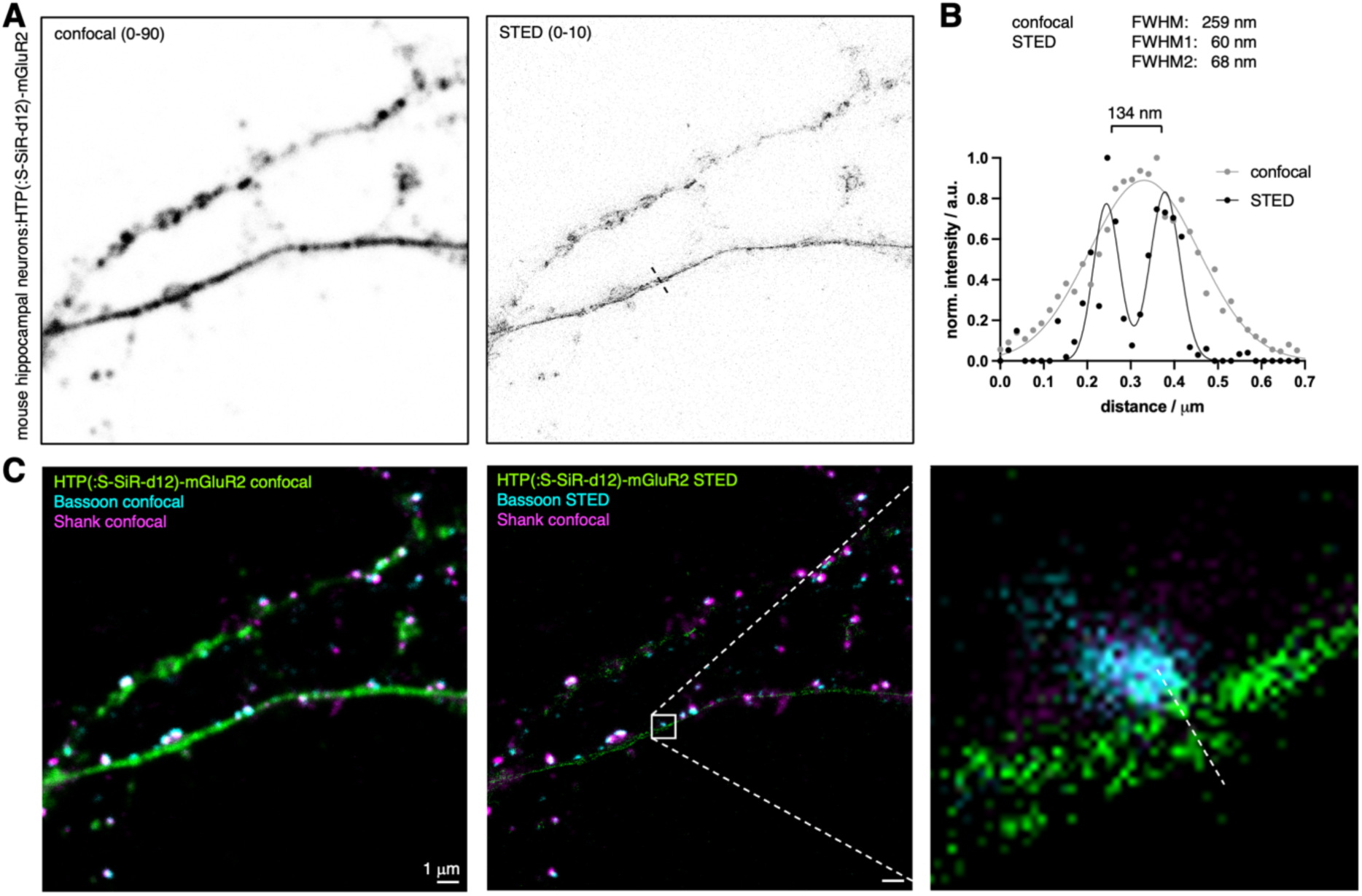
STED super-resolution imaging of surface HTP(:S-SiR-d12)-mGluR2. **A)** Confocal and STED images of HTP(:S-SiR-d12)-mGluR2 transduced neurons. **B**) Line scan profile of a process comparing confocal to STED performance, yielding a resolution of 134 nm across the ultrastructure. **C**) Confocal and dual color STED with zoom in of the process reported in (B).

ATTO 647N is a prominent dye that has been employed in STED^24,25^ and structured illumination microscopy (SIM)^26,27^ super-resolution imaging, and its HTL derivative has been used to stain neurons in *Drosophila*^28^. It is characterized to be a bright dye in the far-red, which is favorable. However, when examining its molecular structure, we noted a four carbon linker on the 3 position for conjugation (**Fig. 5A**), which could exacerbate its intrinsic stickyness^29^. Moreover, we suspected that this linker would slow binding to HTP, given that secondary interaction sites between the HTP surface and rhodamine dyes are known to underlie the rapid labelling speed of dye-HTL conjugates, whereas ligands that do not bear a properly positioned dye or lack a dye are known to bind HTP inefficiently^30^. This limitation has been recently overcome by the design of a second-generation HTL.2^31^, which dramatically improves the efficiency of the DART (drug acutely restricted by tethering) approach^32^.

**Figure 5:**
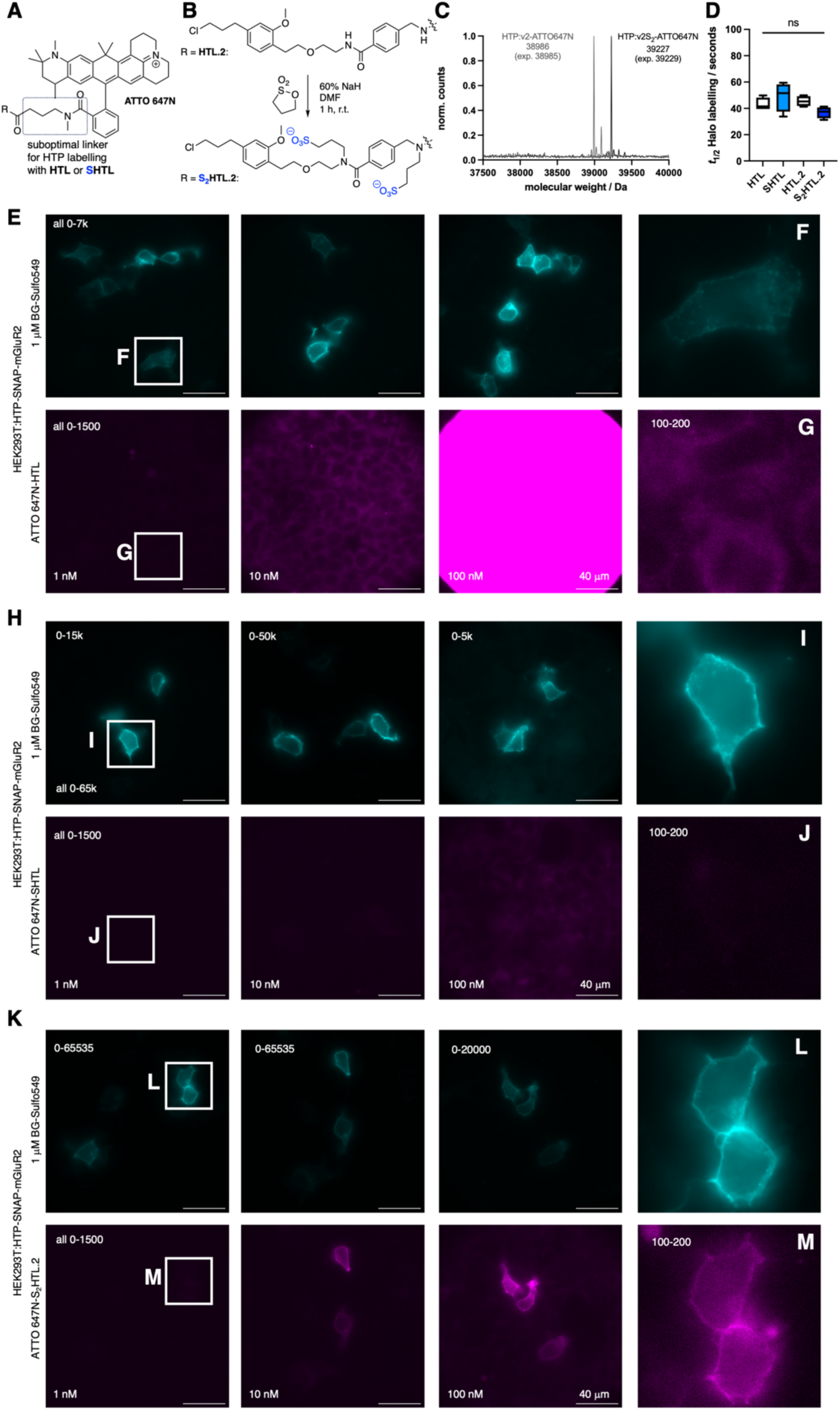
Sulfonation on the HaloTag ligand v2.0 (HTL.2) for improved labelling with sticky ATTO 647N. **A)** Chemical structure of ATTO 647N, with a *N*-methyl amidated additional four carbon linker on the 3 position, which disallows proper dye:HTP secondary interactions. **B**) Sulfonation protocol on second version HaloTag ligand HTL.2 yields double sulfonated S_2_HTL.2. **C**) qTOF full protein mass spectrometry of recombinant HTP labelled with ATTO 647N-HTL.2 and ATTO 647N-S_2_HTL.2. **E**) HTP-SNAP-mGluR2 transfected HEK293 cells, labelled with BG-Sulfo549 (1 uM) and different concentrations of ATTO 647N-HTL.2 for 10 minutes prior to fixation and imaging gives rise to unspecific signal. **F, G**) Zoom ins and brightness contrast adjusted images from (E). **H-J**) As for (E-G) but with different concentrations of ATTO 647N-SHTL leads to image improvements by removing unspecific signals. **K-M**) As for (E-G) but with different concentrations of ATTO 647N-S_2_HTL.2 allows clear membrane labelling even at 1 nM.

To examine whether our sulfonation approach could extend to HTL.2, we synthesized ATTO 647N-HTL and ATTO 647N-HTL.2, before subjecting both to the sulfonation protocol to yield ATTO 647N-SHTL and ATTO 647N-S_2_HTL.2, the latter of which bears two sulfonates as both amides are alkylated (**Fig. 5B**). To characterize nonspecific stickiness, we incubated non-transfected HEK293 cells with 1000 nM of each compound, observing massive non-specific staining for ATTO 647N-HTL (**Fig. S7B**), which was reduced for ATTO 647N-SHTL and ATTO 647N-HTL.2 (**Fig. S7C, D**), and lowest for ATTO 647N-S_2_HTL.2 (**Fig. S7E**).

We confirmed that the ATTO 647-(S)HTL (**Fig. S7A**) and ATTO 647-(S_2_)HTL.2 (**Fig. 5C**) cleanly react with recombinant HTP. We also investigated labelling kinetics by incubating 50 nM solutions of the HTL ligands with an excess of recombinant HTP (200 nM) while tracing fluorescence polarization. While this did not reveal detectible differences (**Fig. 5D**), the assay lacked the sensitivity needed to examine the biologically relevant case, in which low-nanomolar HTL in free solution outnumbers immobilized HTP molecules. We thus turned to the application of interest, using HEK293 cells encoding HTP-SNAP-mGluR2, incubated with HTL ligands in a titration series (1, 10 and 100 nM) for only 10 minutes before fixation. We used sparse expression with low amounts of DNA (50 ng *vs.* 400 ng as in Fig. 2) to ensure that the labelling reaction does not deplete ligand in free solution, particularly at the low concentrations. We co-labelled with the cell-impermeable BG-SulfoJF_549_ (1 uM) to visualize the ideal pattern of surface-only labelling (note: HTP and SNAP are both extracellular).

The non-sulfonated ATTO 647N-HTL exhibited little labelling at 1 to 10 nM, while 100 to 1000 nM yielded predominantly nonspecific intracellular labelling that overshadowed any surface labelling (**Fig. 5E-G**). We believe this represents nonspecific stickiness as it was indistinguishable to that seen in mock-transfected cells, lacking HTP. The sulfonated first-generation ligand, ATTO 647N-SHTL, substantially reduced this nonspecific intracellular labelling in mock cells (**Fig. 5H-J**), however surface-specific HTP labelling in transfected cells was inefficient, appearing incomplete even at 100 nM; thus, surface labelling could be seen, but with bothersome nonspecific labelling. By contrast, the sulfonated second-generation ATTO 647N-S_2_HTL.2 achieved clear labelling when delivered at only 100 nM, revealing bright surface labelling with negligible background (**Fig. 5K**). Even with 1 nM, clean surface labelling was evident after adjusting brightness and contrast (**Fig. 5L, M**). The non-sulfonated ATTO 647N-HTL.2 also exhibited efficient labelling (**Fig. S7F-H**), however, nonspecific intracellular was seen if applied at 1000 nM. Thus, by combining HTL.2 with sulfonation, ATTO 647N-S_2_HTL.2 performed better than all other ligands tested.

The power of chemical biology is often limited by reagent availability. While popular probes are commercially available, variants developed in an academic setting are not always readily available, particularly over longer periods or in larger quantities. We decided to tackle this issue by testing if our one-step protocol to make dye-HTL reagents impermeable can be performed on a small scale without the need for purification, so that any laboratory can perform the synthesis on demand. To facilitate this, we switched the base to sodium *tert*-butoxide, which is soluble to 100 mM in DMSO, making it easier to handle than sodium hydride (**Fig. 6A**). Neat 1,3-propane sultone was warmed to 35 °C (melting point 32 °C) to obtain a liquid that can straightforwardly be pipetted (**Fig. 6A**). We tested a quick protocol, for which 5 nmol of a dye-HTL is dissolved in 4 μL of base solution, before addition of 1 μL of the sultone. Incubation for 5 minutes at room temperature before quenching with 5 μL PBS yields a 0.5 mM dye-SHTL solution that can be directly used for cell application and imaging (**Fig. 6B**). We tested this protocol on commercially available HTL conjugates of JF_549_ and JF_646_, as well as HTL conjugates of our in-house developed probes TMR-d12 and SiR-d12 (**Fig. 6C**). Reactions were >99% quantitative according to LCMS analysis (**Fig. 6D**). It should be noted that the PBS quenching step hydrolyzes the remaining sultone to an impermeable sulfonate, and neutralizes the base. Importantly, we observed no toxicity within 30 minutes of the reaction cocktail when incubating HEK293 cells over 30 min with a 1:1000-dilution (resulting in: [DMSO] = 564 μM or 0.1%; [sultone] = 108 μM; [KOtBu] = 4 μM) tested with a propidium iodide assay for cell death (**Fig. S8**). Furthermore, after exposing HEK293 cells to this cocktail for 24 hours, no effect on cell viability was observed by a WST-1 assay that measures metabolic impact (**Fig. S9**). Higher concentrations led to cell death (LD_50_ = 2.8 mM DMSO / 535 μM sultone / 20 μM KO*t*Bu), presumably caused by *tert*-butanol (i.p. LD_50_ (mouse) = 399 mg/kg), supported by the fact that 14.1 mM DMSO concentrations did not affect viability, and no toxicity reports exist for 3-hydroxypropane-1-sulfonic acid in the National Library of Medicine. Therefore, using a 1:10,000 dilution of the freshly prepared TMR-d12-SHTL (i.e. 50 nM) together with BG-Sulfo646 on HEK293:SNAP-HTP-mGluR2 (both tags on the extracellular side fused to metabotropic glutamate receptor 2) expressing cells (**Fig. 6E**) in confocal imaging, and observed clear co-localization of both channels when using TMR-d12-SHTL and BG-Sulfo646. Similar performance was observed for JF_549_-SHTL, SiR-d12-SHTL, and JF_646_-SHTL (**Fig. 6F**), with the latter showing the cleanest performance.

**Figure 6:**
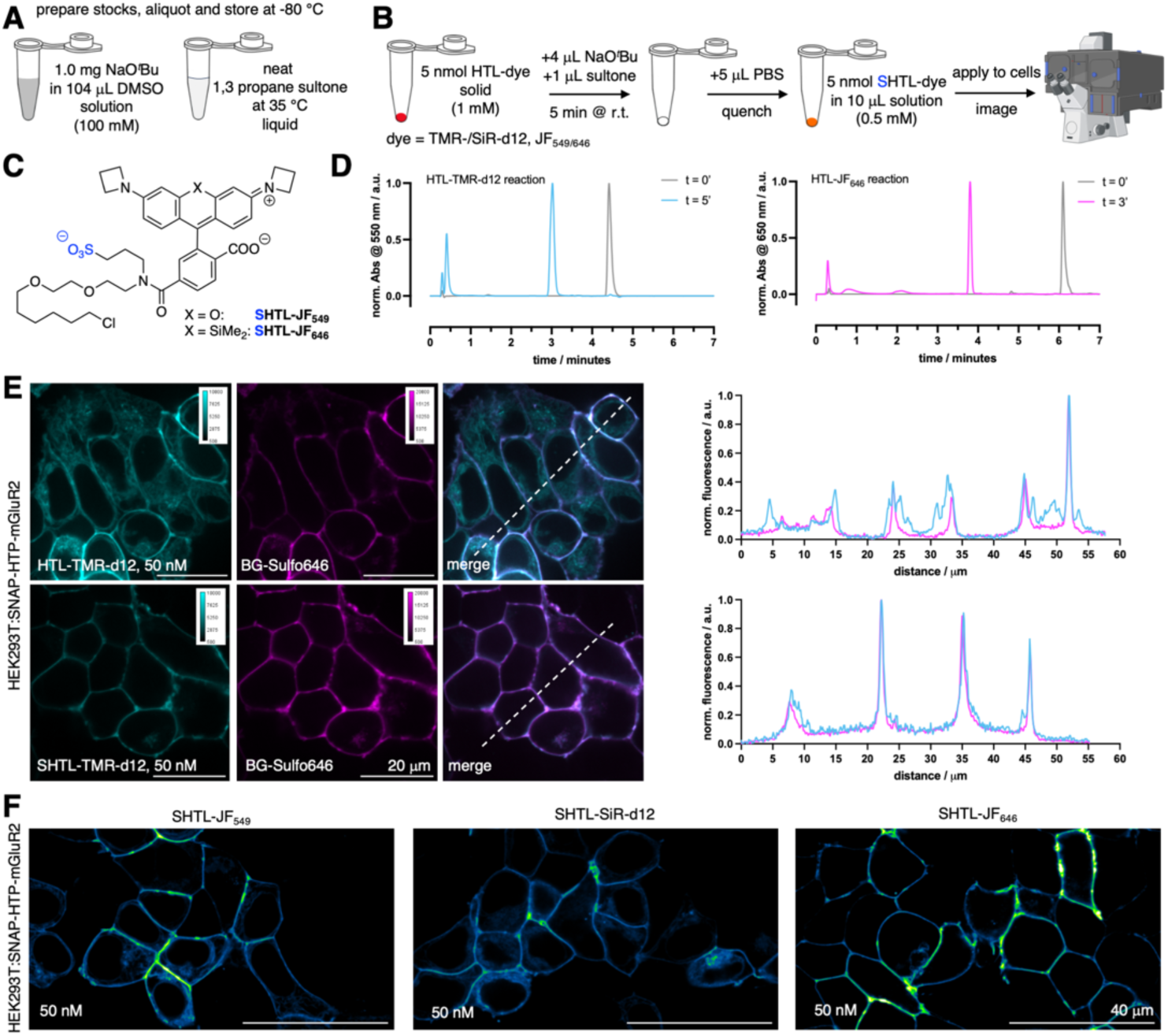
One step protocol on small scale to synthesize and apply dye-SHTL. **A)** Required stock solutions. **B)** Outlined 5-minute synthetic protocol (partly created in biorender.com). **C)** Structures of JF_549/646_-SHTL. **D)** LCMS traces of the reaction for TMR-d12-HTL and JF_646_-HTL. **E)** Confocal imaging of HEK293:SNAP-HTP-mGluR2 transfected cells with TMR-d12-(S)HTL (50 nM) and BG-Sulfo646 (50 nM) including line scans. **F)** As for E, but with JF_549_-SHTL, SiR-d12-SHTL and JF_646_-SHTL.

## DISCUSSION

The development of bright (and impermeable) fluorophores for microscopy has garnered increasing interest,^11,33,34^ particularly in the area of shadow imaging^35,36^ and protein labelling^7,13,37,38^. In this study, we aimed to install a sulfonate on the amide bond of existing dye-HTL substrates to render them impermeable to a cell’s plasma membrane lipid bilayer. This strategy was applied on rhodamine scaffolds, since these dye-HTL molecules only bear one acidic proton, and furthermore, in basic solutions form a non-fluorescent spirolactone form, preventing off-target alkylation on the carboxylate. The majority of dye-HTL reagents should be amenable to this method, however, the approach would not be applicable to certain scaffolds with more possible attachment sites, including biotin, dyes with nucleophilic sites, for instance NH-anilines (e.g. Rho6G, SiR595) or hydroxy groups (e.g. Oregon Green). We observed *cis*- and *trans*-amide mixture in ^1^H NMR (see SI), which we were not able to separate, however, we observed full protein labelling by mass spectrometry using recombinant HTP, and observed no difference in labelling kinetics via fluorescence polarization compared to non-sulfonated HTL substrates. We demonstrate impermeability of SHTL conjugates in live cell staining using widefield microscopy on HEK cells transfected with extra- and intracellular localized SNAP and HTP. This allowed live-cell labelling of HTP-mGluR2 transduced hippocampal neurons with SiR-d12-SHTL prior to fixation, antibody staining and confocal imaging. By either specifically targeting the extracellular, exposed HTP-mGluR2 protein pools with SiR-d12-SHTL, or the total protein pool with SiR-d12-HTL, we found that there is a mix of surface and intracellular populations in the soma and both dendritic and axonal processes. Critically, SiR-d12-SHTL allowed us to find the presence of a surface pool of axonal and presynaptically-localized mGluR2. As the trafficking mechanisms and nanolocalization of mGluRs is a highly active field of study,^20,22,23,39^ our observations point towards many surface and intracellular subpopulations of mGluR2, both in the soma and processes/synapses, which will warrant future study, amenable to super-resolution STED nanoscopy. Although we used an overexpression system, our findings open up interesting avenues to probe GPCRs and other receptors in different cellular compartments with precision for surface versus intracellular pools, which we aim to perform on endogenous HTP-fused proteins in the future. For these reasons, we showcase ATTO 647N on the HTL.2 substrate to exhibit cleaner labelling even at low doses down to 1 nM, which may be essential when applied to complex tissues or in live animals, where high concentrations are difficult to achieve. In addition, ATTO 647N has been reported be sticky, which may lead to nonspecific background labeling,^40,41^ and thus the addition of two negative charges on S_2_HTL.2 offers even greater attenuation of nonspecific stickiness. Other dyes that may be interesting, especially due to their performance in the near-infrared, include 2XR-1 (ref^42^) and heptamethine cyanines^43^. Finally and generally, a one-step protocol without means of purification for direct application in biology are attractive,^44,45^ and we have added to this portfolio. Although the reaction is near-quantitative, small amounts of starting material left may give rise to intracellular staining (**Fig. S10**). As such, the use of far-red JF_646_-SHTL alongside with 10,000-fold dilutions of the reaction mixture are recommended for surface labelling. We provide a one-page step-by-step protocol for straightforward implementation in the Supporting Information.

## SUMMARY

We report on the synthesis and application of sulfonated HaloTag substrates, which can be performed in one step, to label HTP-fused cell surface proteins in the red and far-red. Demonstrated in live cell widefield imaging in HEK293 cells, and in confocal and STED super-resolution microscopy on HTP-mGluR2 transduced hippocampal neurons, we confirm their impermeability and the ability to precisely localize the surface population of a GPCR involved in neurotransmission. We expand the strategy to the second-generation HTL.2, and provide a simple protocol that any laboratory can perform on small scale without the need for purification. While not all substrates are amenable to this approach, we envision wide adoption.

## AUTHOR CONTRIBUTION

Conceptualization and Methodology: JB; Formal analysis and investigation: KR, SS, CHO, MK, ET, UP, RB, NL and JB; Writing –Original Draft: JL and JB; Reviewing and Editing: KR, JL, NL, ML, MRT, BCS, JH and JB; Visualization: KR and JB; Supervision: NL and JB; Funding Acquisition: JL, NL and JB.

## ACKNOWLEDGEMENTS

This project has received funding from the European Union’s Horizon Europe Framework Programme (deuterON, grant agreement no. 101042046 to JB). This work was supported by the German Research Foundation Excellence Strategy EXC-2049-390688087 (NL) and CRC 1286 “Quantitative Synaptology” project A11 (NL). HTL.2 reagents were supported by NIH grants 1RF1MH117055 and 1DP2MH1194025 (to MRT). JL is supported by NIH grant R01NS129904.

## MATERIALS AND METHODS

Chemical synthesis and characterization, modelling, measurements of photophysical and kinetic parameters, and procedures in cell culture, molecular biology and imaging are reported in the Supporting Information.

