## Supplementary Information for "A one-step protocol to generate impermeable fluorescent HaloTag substrates for *in situ* live cell application and super-resolution imaging"

[1] Leibniz-Forschungsinstitut für Molekulare Pharmakologie (FMP), 13125 Berlin, Germany.
[2] Weill Cornell Medicine, Department of Biochemistry, NY, USA.
[3] Duke University, Department of Biomedical Engineering, Durham, North Carolina 27708, USA
[4] Duke University, Department of Chemistry, Durham, North Carolina 27708, USA

* Correspondence should be addressed to:


### 1 General

All chemical reagents and anhydrous solvents for synthesis were purchased from commercial suppliers (Sigma-Aldrich, Roth) and were used without further purification if not stated otherwise.

NMR spectra were recorded at 300 K in deuterated solvents on a Bruker AV-III spectrometer using a room- temperature 5 mm broadband probe equipped with one-axis self-shielded gradients and calibrated to residual solvent peaks ($^{1}H$/$^{13}C$ in ppm): $D_2O$ (4.65/N.A.). Multiplicities are abbreviated as follows: s = singlet, d = doublet, t = triplet, q = quartet, p = pentet, h = heptet, br = broad, m = multiplet. Coupling constants *J* are reported in Hz. Spectra are reported based on appearance, not on theoretical multiplicities derived from structural information.

UPLC-UV/Vis for purity assessment was performed on an Agilent 1260 Infinity II LC System equipped with Agilent SB-C18 column (1.8 µm, 2.1 × 50 mm). Buffer A: 0.1% FA in $H_2O$ Buffer B: 0.1% FA acetonitrile. The typical gradient was from 10% B for 1.0 min → gradient to 95% B over 5 min → 95% B for 1.0 min with 0.6 mL/min flow or from 30% B for 1.0 min → gradient to 95% B over 5 min. For ATTO 647N-HTL, ATTO 647N-SHTL and ATTO 647-$S_2$HTL.2 50% B for 1.0 min → gradient to 95% B over 5 min was used and for ATTO 647-HTL.2 70% B for 1.0 min → gradient to 95% B over 5 min was used instead. Retention times ($t_R$) are given in minutes (min). Chromatograms were imported into Graphpad Prism8 and purity was determined by calculating AUC ratios.

Preparative or semi-preparative HPLC was performed on an Agilent 1260 Infinity II LC System equipped with columns as followed: preparative column –Reprospher 100 C18 columns (10 µm: 50 x 30 mm at 20 mL/min flow rate; semi-preparative column – 5 µm: 250 x 10 mm at 4 mL/min flow rate. Eluents A (0.1% TFA in $H_2O$) and B (0.1% TFA in MeCN) were applied as a linear gradient. Peak detection was performed at maximal absorbance wavelength.

For HRMS, samples were analyzed on Orbitrap Fusion mass spectrometer (Thermo Fisher Scientific). MS scans were acquired in a range of 350 to 1500 m/z. MS1 scans were acquired in the Orbitrap with a mass resolution of 120,000 with an AGC target value of 4e5 and 50 ms injection time. MS2 scans were acquired in the ion trap with an AGC target value of 1e4 and 35 ms injection time. Precursor ions with charge states 2-4 were isolated with an isolation window of 1.6 m/z and 40 sec dynamic exclusion. Precursor ions were fragmented using higher-energy collisional dissociation (HCD) with 30% normalized collision energy.

#### 2 Cell lines and microscopy

##### 2.1 Culture, transfection and staining

HEK293T cells were cultured in growth media (DMEM, Glutamax, 4.5 g Glucose, 10% FCS, 1% PS; Invitrogen) at 37 °C and 5% $CO_2$. 50 000 cells per well were seeded on 8-well µL slides (Ibidi) previously coated with poly-L-lysine (Aldrich, mol wt 70 000–150 000). The next day, 400 ng DNA was transfected using 0.8 µL Jet Prime reagent in 40 µL Jet Prime buffer (VWR) per well/plasmid. Media was exchanged against antibiotic-free media before the transfection mix was pipetted on the cells. After 4 hours incubation at 37 °C and 5% $CO_2$, medium was exchanged against growth media, and after an additional 24 hours, cells were stained. All dyes for widefield imaging were used at a concentration of 500 nM with the addition of Hoechst 33342 at 1 µM, in growth media. Cells were stained at 37 °C and 5% $CO_2$ for 30 minutes. Afterwards cells were washed once in growth media and imaged in fluorobrite (Invitrogen). For confocal microscopy all dyes were additionally used at a concentration of 50 nM.

##### 2.2 Live Cell Widefield Imaging

Living cells were imaged in fluorobrite (Invitrogen) using an epifluorescence Nikon Ti-E microscope, equipped with pE4000 (cool LED), Penta Cube (AHF 66-615), 60× oil NA 1.49 (Apo TIRF Nikon) and imaged on a sCMOS camera (Prime 95B, Photometrics) operated by NIS Elements (Nikon). For excitation the following LED wavelengths were used: Hoechst – 405 nm, $JF_{549}$, TMR-d12 – 550 nm, $JF_{646}$, SiR-d12 – 635 nm.
Line scans were drawn with the polygon selection in ImageJ and the grey values obtained and plotted against the distance in Graphpad Prism 10, in which values were also normalized.

##### 2.3 FLIM

Drops of 10-20 ul of 2-5 µM Fluorophor where spotted in u-Slide 8 Well Glass Bootm (Ibidi #80877). Fluorescence lifetime was measured on a Leica SP8 TCS STED FALCON (Leica Microsystems) equipped with a pulsed white-light excitation laser (80 MHz repetition rate, NKT Photonics), a 100× objective (HC PL APO CS2 100×/1.40 NA oil), a temperature controlled chamber at room temperature, operated by LAS X. A Hybrid detector produces FLIM images of 512 × 512 pxl with 113 nm per pxl after 10 frame repetitions. Solution of 5 µM Fluorescein was used as a reference, excited at 488nm and the lifetime of the collected em from λ = 503-580 nm was 3.8 ns. HTL and TMR-d12-SHTL solutions (2 µM) were excited using λ = 561 nm, emission signals were captured at λ = 577–650 nm before and after addition of 10 µM purified HaloTag protein. HTL and SiR-d12-SHTL were excited using λ = 640 nm, emission signals were captured at λ = 655–750 nm. Fluorescence lifetime decay curves from selected regions were fitted with one exponential function and the lifetime is reported for each region.

##### 2.4 Hippocampal neurons

Primary hippocampal neurons were cultured as previously described in ref[1]. Briefly, hippocampi were dissected from wild-type C57bl/6J P0-P1 mice, and were incubated in 37 °C gently shaking in a solution containing 0.2 mg/ml L-cysteine, 1 mM $CaCl_2$, 0.5 mM ethylenediaminetetraacetic acid (EDTA) and 25 units/ml of papain (Worthington Biochemicals), pH 8 in Dulbecco's modified eagle medium. After one hour, the solution was replaced by a prewarmed solution containing 2.5 mg/ml Bovine Serum Albumin, 2.5 mg/ml trypsin inhibitor (Sigma-Aldrich T9253), 1% Fetal Bovine Serum, heat inactivated (FBS; e.g., Gibco™ A15-104) in Dulbecco's modified eagle medium, and the hippocampi are incubated at

37 °C gently shaking. The solution is then replaced by neuronal culture medium (Neurobasal™-A medium supplemented with 2% B-27™ Plus Supplement, 1% GlutaMAX™ supplement, and 1% Penicillin-Streptomycin), and the hippocampi are gently triturated to produce a cell suspension that is then plated on PLL-coated coverslips for 2-3 weeks at 37 °C and 5% $CO_2$ before experiments begin. The culture contains primarily neurons and few astrocytes, and is thus ideal for imaging.

At day-in-vitro 1-2, neurons were infected with lentiviral particles encoding for HTP-mGluR2, prepared according to the protocol published in ref[2] and modified as in ref[3], using a modified version of the Addgene plasmids #8454 and #8455 (ref[4]) and under the synapsin promoter.

At day-in-vitro 14-17, neurons were incubated for 30 minutes with the indicated dyes at 37°C and 5% $CO_2$, washed two times in PBS to remove unbound dyes and cell debris, fixed with cold 1% paraformaldehyde (PFA) in PBS for 10 minutes, and then washed two times in PBS to remove the PFA.

The cells were permeabilized with cold 0.25% Triton-X-100 in PBS for 10 minutes and rinsed once in pure PBS. To block nonspecific binding sites, the cells were incubated in cold 0.3% NGS in PBS for 20 minutes. Next, neurons were stained with the following antibodies for 1 hour at RT: Guinea pig polyclonal Shank 2 (Synaptic Systems 162 204, diluted 1:250), Chicken polyclonal MAP2 (Novus Biologicals NB300-213, diluted 1:1000), and Mouse monoclonal anti Bassoon (Abcam, AB_82958, diluted 1:250), in PBS containing 0.3% NGS. To remove unbound antibodies, the cultures were rinsed three times with blocking solution before applying the following secondary antibodies conjugated with fluorophores: Anti-Chicken-AF405 (Abcam AB_175674), Anti-Guinea Pig-CF488 (Biotium CF® AB_20169-1) and Anti-Mouse-AF594 (Invitrogen A32744). The secondary antibodies were diluted 1:500. The neurons were incubated with the secondary antibodies for 45 minutes at RT in the dark and washed eight times in PBS to remove unbound products. Next, neurons were fixed with 4% PFA in PBS for 15 minutes at RT and then washed two times in PBS to remove excessive PFA. The coverslips were rinsed once by dipping shortly into dd$H_2O$, mounted with ProLong™ Gold Antifade Mountant (Invitrogen™ P36934) on microscope glass slides, and stored in the dark at RT for 48 hours to dry before imaging.

#### 2.5 Confocal Imaging

Confocal microscopy on transduced neurons was performed using a Leica SP8 TCS STED FALCON (Leica Microsystems) equipped with a pulsed white-light excitation laser (80 MHz repetition rate, NKT Photonics), a 100x objective (HC PL APO CS2 100×/1.40 NA oil), a temperature-controlled chamber and operated by LAS X. SiR-d12 was excited using $\lambda$ = 647 nm and emission signals were captured at $\lambda$ = 656-751 nm. Confocal images were collected using a time gated Hybrid detector (0.5–6 ns). Images of 1024 x 1024 pixel had a pixel size of 113.64 nm. Regions of interest were manually drawn using FIJI, and mean gray values plotted in Graphpad Prism 10. Statistics were calculated in Graphpad Prism 10.

### 3 Quantum Yield

Absolute quantum yields were taken at a steady-state UV-Vis absorption spectroscopy on a Hamamatsu PL Quantum Yield Spectrometer C11347 in 1 cm quartz cuvettes. The solutions were prepared to have an absorbance between 0.05 and 0.1 at 549 nm for TMR in PBS and at 652 nm for SiR in PBS and quantum yields were determined on the same instrument equipped with an integrating sphere.

### 4 Synthesis

#### 4.1 General procedure A for SHTL conjugates

A 5 nmol aliquot of HTL conjugated fluorophore was dissolved in 4 µL *tert*-butoxide (100 mM in DMSO) and 1 µL of 35 °C warm 1,3-propane sultone was added. After the reaction was complete (monitored by LCMS, which takes around 5 min), the reaction was quenched with 5 µL PBS and used without further purification.

##### 4.2 3-(3-(6-(Bis(methyl-d$_3$)amino)-3-(bis(methyl-d$_3$)iminio)-3*H*-xanthen-9-yl)-4-carboxy-*N*-(2-(2-((6-chlorohexyl)oxy)ethoxy)ethyl)benzamido)propane-1-sulfonate (TMR-d12-SHTL)

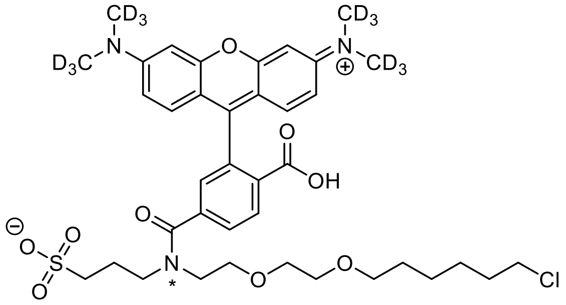

**TMR-d12-SHTL** was prepared according to general procedure A with TMR-d12-HTL or by the following procedure:

TMR-d12-HTL (1.88 mg, 2.90 µmol, 1.0 equiv.) was dissolved in DMF, before NaH 60% in mineral oil (2.32 mg, 58.0 µmol, 20.0 equiv.) and 1,3-propane sultone (1.77 mg, 14.5 µmol, 5.0 equiv.) were added. The reaction was stirred for 1 h at rt, before it was quenched by addition of glacial HOAc (50 µL). The reaction mixture was subjected to RP-HPLC to obtain 2.10 mg (2.73 µmol) of the desired compound as a blue powder in 94% yield. NMR spectra are reported for *cis-*/*trans*-amide isomers.

**$^1$H NMR** (600 MHz, DMSO-d$_6$) $\delta$ [ppm] = 8.25 (q, $J$ = 7.7 Hz, 2H), 7.79 (d, $J$ = 7.9 Hz, 1H), 7.73 (d, $J$ = 7.9 Hz, 1H), 7.52 (s, 1H), 7.38 (s, 1H), 7.17 (s, 4H), 7.10 (s, 4H), 6.95 (s, 4H), 3.58 (t, $J$ = 8.5 Hz, 8H), 3.51 (q, $J$ = 6.6 Hz, 10H), 3.46 (d, $J$ = 3.3 Hz, 2H), 3.20 (s, 2H), 3.14 (t, $J$ = 6.5 Hz, 2H), 2.55 (s, 4H), 2.43 (t, $J$ = 7.3 Hz, 2H), 2.26 (t, $J$ = 7.0 Hz, 2H), 1.88 (t, $J$ = 7.1 Hz, 2H), 1.72 (t, $J$ = 6.8 Hz, 2H), 1.65 (m, $J$ = 7.7 Hz, 4H), 1.42 (q, $J$ = 6.8 Hz, 2H), 1.30 (m, $J$ = 8.0 Hz, 8H), 1.17 (t, $J$ = 7.2 Hz, 2H).

**HRMS** (ESI): calc. for $C_{38}H_{37}D_{12}ClN_3O_9S$ $[M]^+$: 770.3626, found: 770.3632.

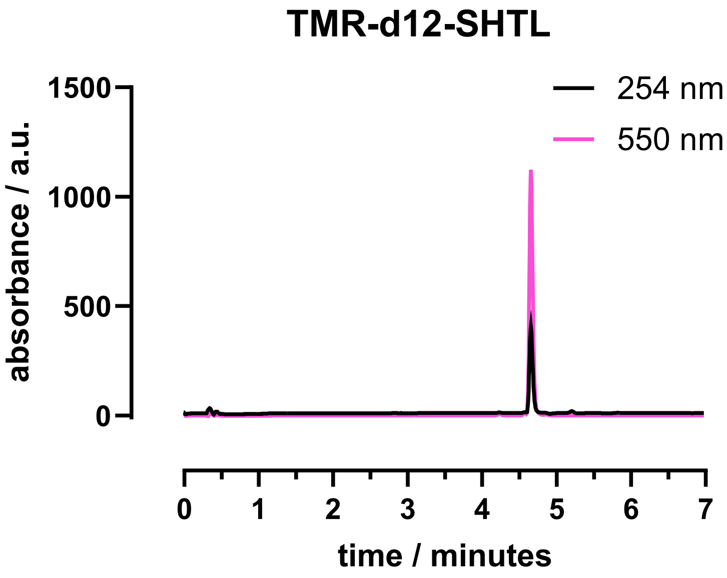

##### 4.3 3-(3-(7-(Bis(methyl-d$_3$)amino)-3-(bis(methyl-d$_3$)iminio)-5,5-dimethyl-3,5-dihydrodibenzo[*b,e*]silin-10-yl)-4-carboxy-*N*-(2-(2-((6-chlorohexyl)oxy)ethoxy)ethyl)benzamido)propane-1-sulfonate (SiR-d12-HTL)

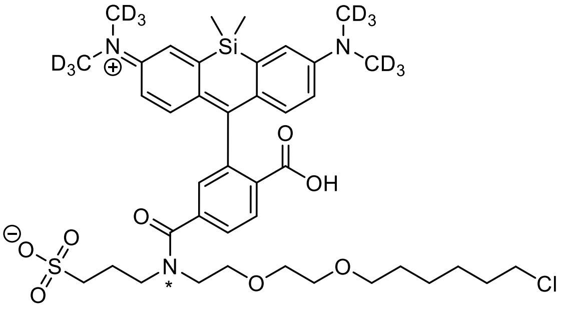

**SiR-d12-SHTL** was prepared according to general procedure A with SiR-d12-HTL or by the following procedure:

SiR-d12-HTL (2.0 mg, 2.90 µmol, 1.0 equiv.) was dissolved in 500 µL DMF, before NaH 60% in mineral oil (2.32 mg, 58.0 µmol, 20.0 equiv.) and 1,3-propane sultone (1.77 mg, 14.5 µmol, 5.0 equiv.) were added. The reaction was stirred for 1 h at rt, before it was quenched by addition of glacial HOAc (50 µL). The reaction mixture was subjected to RP-HPLC to obtain 2.14 mg (2.64 µmol) of the desired compound as a blue powder in 91% yield. NMR spectra are reported for *cis-*/*trans*-amide isomers.

**$^1$H NMR** (600 MHz, $D_2O$) δ [ppm] = 8.18 (dd, *J* = 8.0, 15.1 Hz, 1H), 8.00 (t, *J* = 2.8 Hz, 3H), 7.83 (d, *J* = 8.0 Hz, 1H), 7.79 (d, *J* = 8.0 Hz, 1H), 7.69 (s, 1H), 7.56 (dd, *J* = 2.5, 8.8 Hz, 1H), 7.49 (d, *J* = 8.8 Hz, 1H), 7.40 (s, 1H), 7.32 (d, *J* = 8.9 Hz, 1H), 7.22 (d, *J* = 8.9 Hz, 1H), 3.79-3.71 (m, 1H), 3.66-3.60 (m, 1H), 3.53 (bs, 1H), 3.40 (t, *J* = 7.7 Hz, 1H), 3.37 (t, *J* = 6.6 Hz, 1H), 3.31-3.25 (m, 1H), 2.95 (t, *J* = 7.6 Hz, 1H), 2.87 (t, *J* = 6.4 Hz, 1H), 2.48 (t, *J* = 7.5 Hz, 1H), 2.09 (p, *J* = 7.4 Hz, 1H), 1.85 (p, *J* = 7.3 Hz, 1H), 1.42 (p, *J* = 7.3 Hz, 1H), 1.34 (p, *J* = 7.2 Hz, 1H), 1.17 (p, *J* = 7.2 Hz, 1H), 1.0-0.95 (m, 1H), 0.91 (p, *J* = 7.6 Hz, 1H), 0.84 (p, *J* = 7.6 Hz, 1H), 0.77 (s, 1H), 0.65 (s, 1H), 0.63 (s, 1H).

**HRMS** (ESI): calc. for $C_{40}H_{43}D_{12}ClN_3O_8SSi$ $[M]^+$: 812.3915, found: 812.3915.

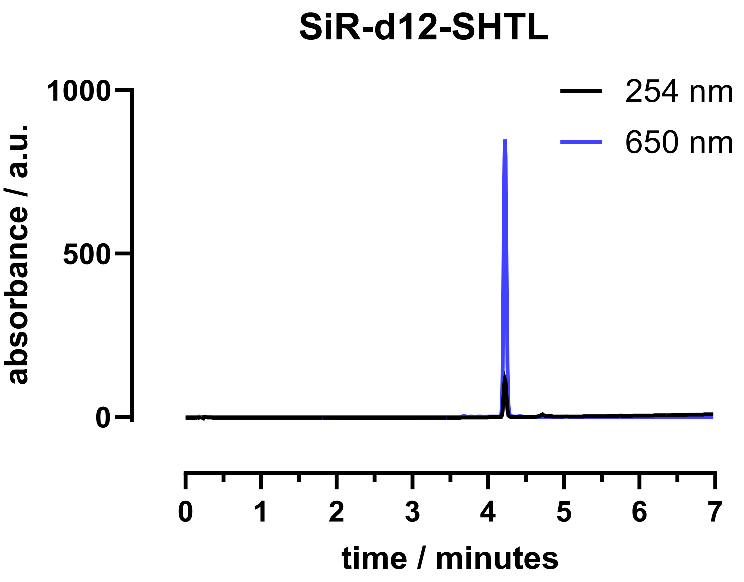

#### 4.4 3-(3-(3-(Azetidin-1-ium-1-ylidene)-6-(azetidin-1-yl)-3*H*-xanthen-9-yl)-4-carboxy-*N*-(2-(2-((6-chlorohexyl)oxy)ethoxy)ethyl)benzamido)propane-1-sulfonate ($JF_{549}$-SHTL)

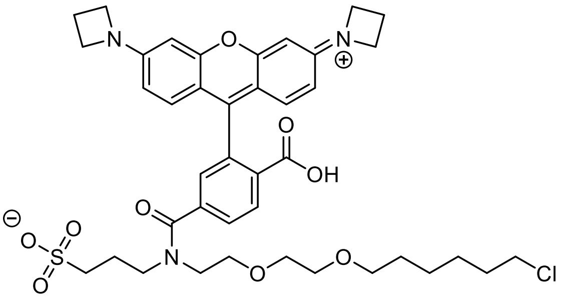

**$JF_{549}$-SHTL** was prepared according to general procedure A with $JF_{549}$-HTL and used without further purification.

**HRMS** (ESI): calc. for $C_{40}H_{49}ClN_3O_9S$ $[M]^+$: 782.2873, found: 782.2873.

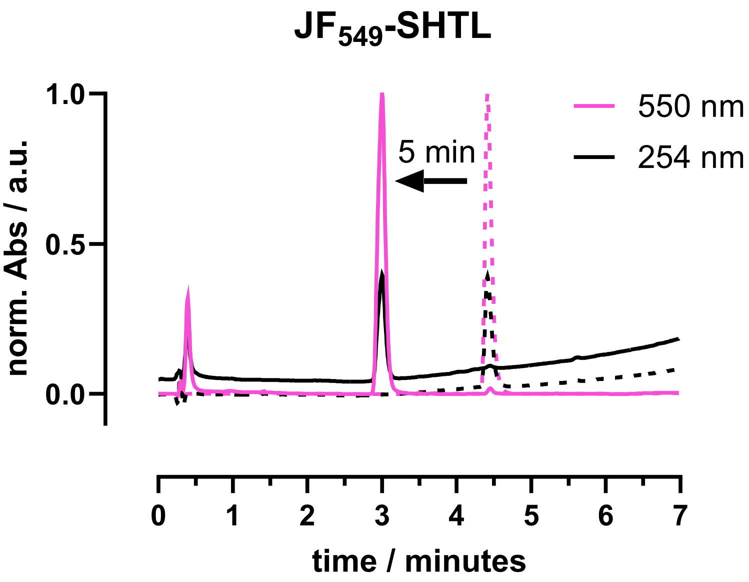

#### 4.5 3-(3-(3-(Azetidin-1-ium-1-ylidene)-7-(azetidin-1-yl)-5,5-dimethyl-3,5-dihydrodibenzo[*b,e*]silin-10-yl)-4-carboxy-*N*-(2-(2-((6-chlorohexyl)oxy)ethoxy)ethyl)benzamido)propane-1-sulfonate ($JF_{646}$-SHTL)

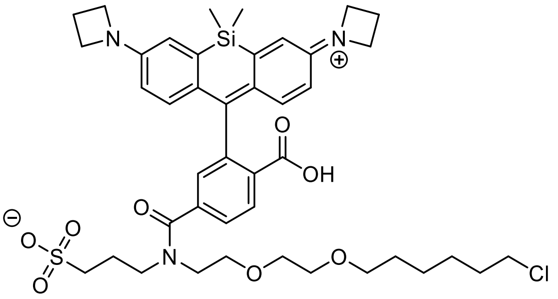

**$JF_{646}$-SHTL** was prepared according to general procedure A with $JF_{646}$-HTL and used without further purification.

**HRMS** (ESI): calc. for $C_{42}H_{55}ClN_3O_8SSi$ $[M]^+$: 824.3162, found: 824.3162.

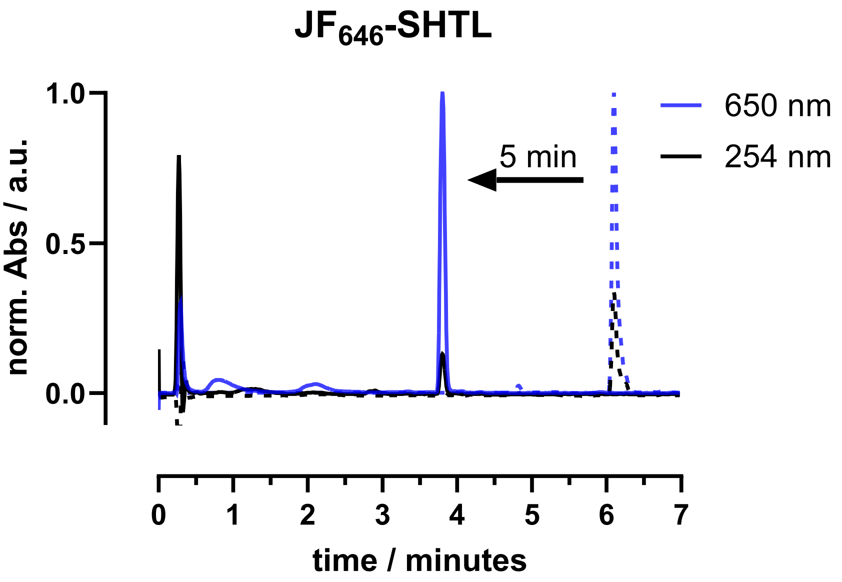

#### 4.6 ATTO 647N-HTL

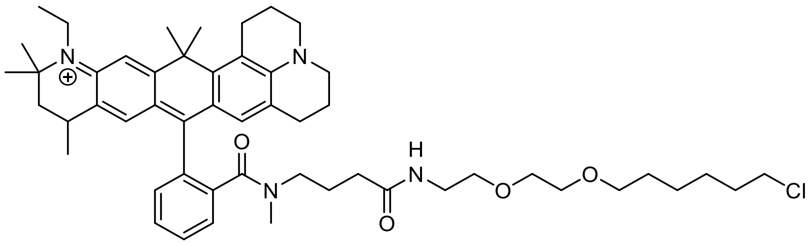

ATTO 647N-NHS (0.5 mg, 673 nmol, 1.0 equiv.) was dissolved in 500 µL DMSO, before DIPEA (0.5 µL, 2.69 µmol, 4.0 equiv.) and 1.2 equiv. HTL-$NH_2$ (0.3 mg, 1.35 µmol, 1.2 equiv.) were added. The mixture was vortexed again and allowed to incubate for 60 min before it was quenched by addition of glacial HOAc (50 µL). HPLC (MeCN:$H_2O$+0.1% TFA = 30:70 to 90:10 over 46 minutes) provided the desired compound (612 nmol, 91%) as blue powder after lyophilization.

**HRMS** (ESI): calc. for $C_{52}H_{72}ClN_4O_4$ $[M]^+$: 851.5237, found: 851.5233.

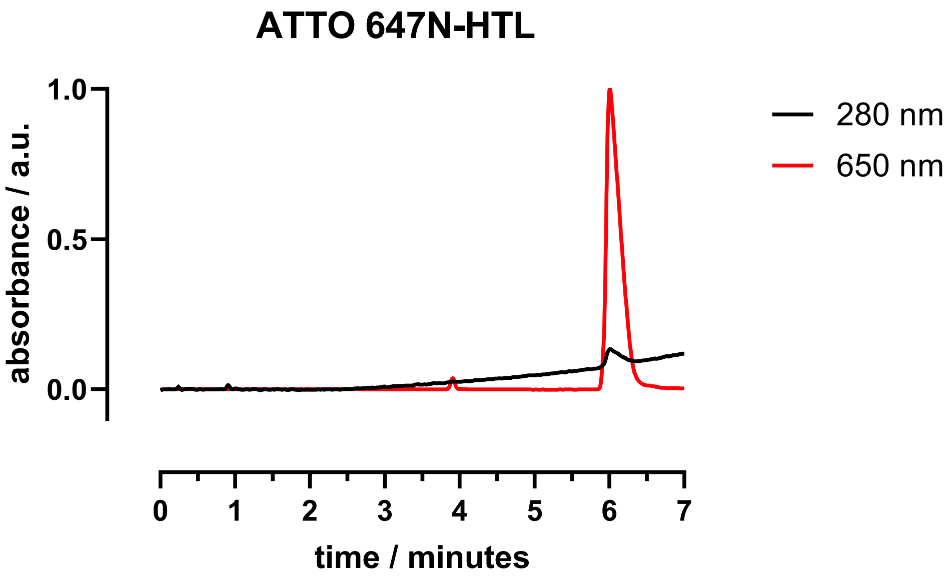

#### 4.7 ATTO 647N-SHTL

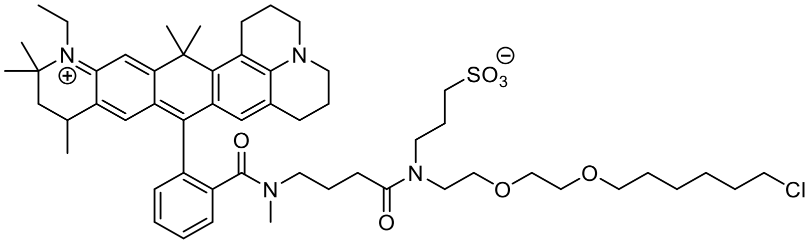

ATTO 647N-HTL (61.0 µg, 72 nmol, 1.0 equiv.) was dissolved in 200 µL DMF, before NaH 60% in mineral oil (57.6 µg, 1.44 µmol, 20.0 equiv.) and 1,3-propane sultone (44 µg, 360 nmol, 5.0 equiv.) was added. The reaction was briefly sonicated, before it was stirred for 6 h at rt. The mixture was quenched by addition of glacial HOAc (50 µL) and subjected to RP-HPLC (MeCN:$H_2O$+0.1% TFA = 30:70 to 90:10 over 46 minutes) to obtain 6.8 µg (7 nmol) of the desired compound as a blue powder in 10% yield, with starting material (40 nmol, 56%) recovered.

**HRMS** (ESI): calc. for $C_{55}H_{78}ClN_4O_7S$ $[M+H]^{2+}$: 487.2674, found: 487.2669.

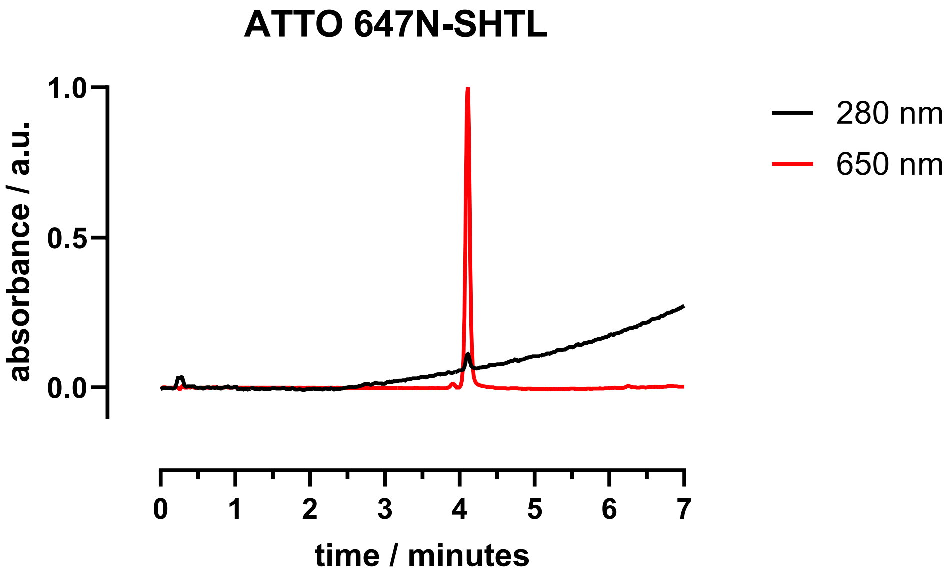

#### 4.8 ATTO 647N-HTL.2

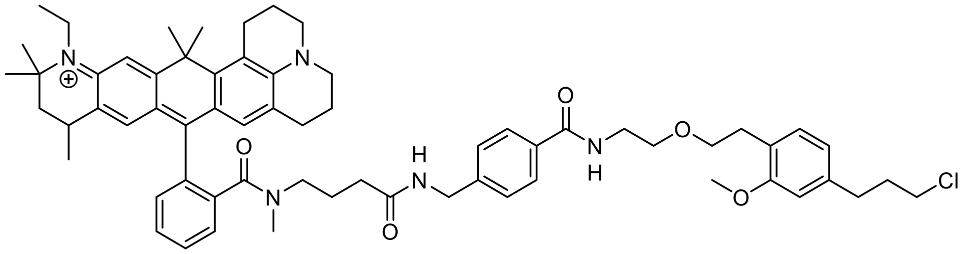

ATTO 647N-NHS (0.5 mg, 673 nmol, 1.0 equiv.) was dissolved in 500 µL DMSO, before DIPEA (0.5 µL, 2.69 µmol, 4.0 equiv.) and HTL.2-$NH_2$ (0.5 mg, 1.35 µmol, 1.2 equiv.) were added. The mixture was vortexed again and allowed to incubate for 60 min before it was quenched by addition of glacial HOAc (50 µL). HPLC (MeCN:$H_2O$+0.1% TFA = 30:70 to 90:10 over 46 minutes) provided the desired compound (599 nmol, 89%) as blue powder after lyophilization.

**HRMS** (ESI): calc. for $C_{64}H_{79}ClN_5O_5$ $[M+H]^{2+}$: 516.7919, found: 516.7909.

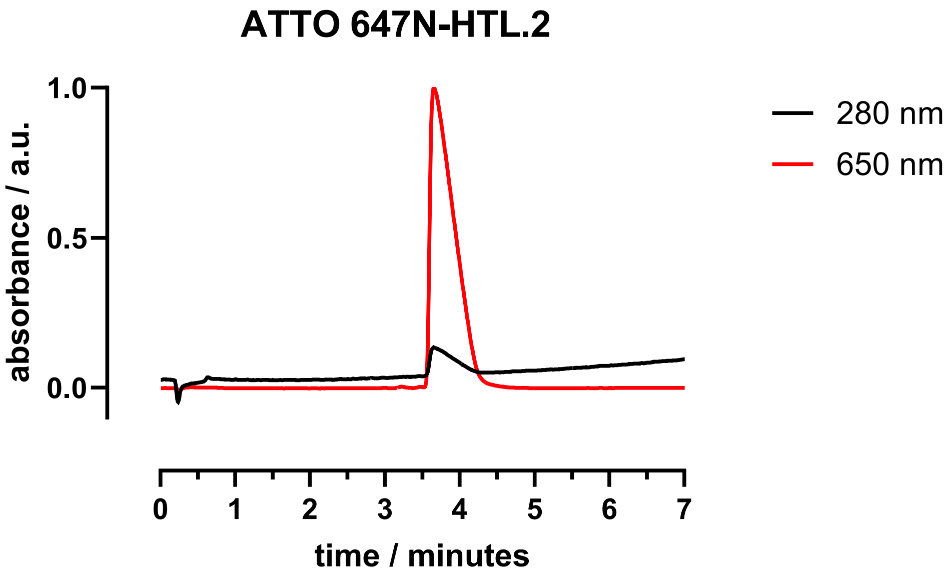

##### 4.9 ATTO 647N-$S_2$HTL.2

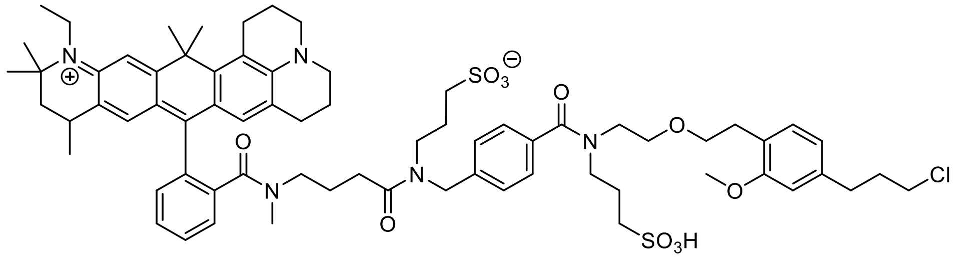

ATTO 647N-HTL.2 (145 µg, 170 nmol, 1.0 equiv.) was dissolved in 200 µL DMF, before NaH 60% in mineral oil (136 µg, 3.4 µmol, 20.0 equiv.) and 1,3-propane sultone (104 µg, 850 nmol, 5.0 equiv.) were added. The reaction was briefly sonicated, before it was stirred for 6 h at rt. The mixture was quenched by addition of glacial HOAc (50 µL) and subjected to RP-HPLC (MeCN:$H_2O$+0.1% TFA = 10:90 to 90:10 over 46 minutes) to obtain 29 µg (23 nmol) of the desired compound as a blue powder in 14% yield, with starting material (68 nmol, 40%) recovered.

**HRMS** (ESI): calc. for $C_{70}H_{91}ClN_5O_{11}S_2$ $[M+H]^{2+}$: 638.7956, found: 638.7967.

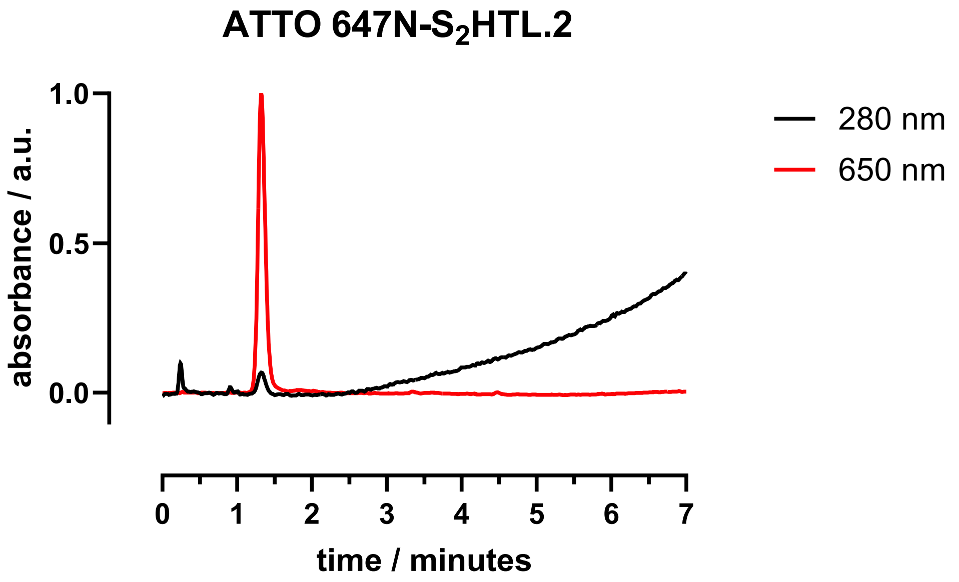

### 5 NMR-Spectra

#### 5.1 3-(3-(6-(Bis(methyl-d3)amino)-3-(bis(methyl-d3)iminio)-3*H*-xanthen-9-yl)-4-carboxy-*N*-(2-(2-((6-chlorohexyl)oxy)ethoxy)ethyl)benzamido)propane-1-sulfonate (TMR-d12-SHTL)

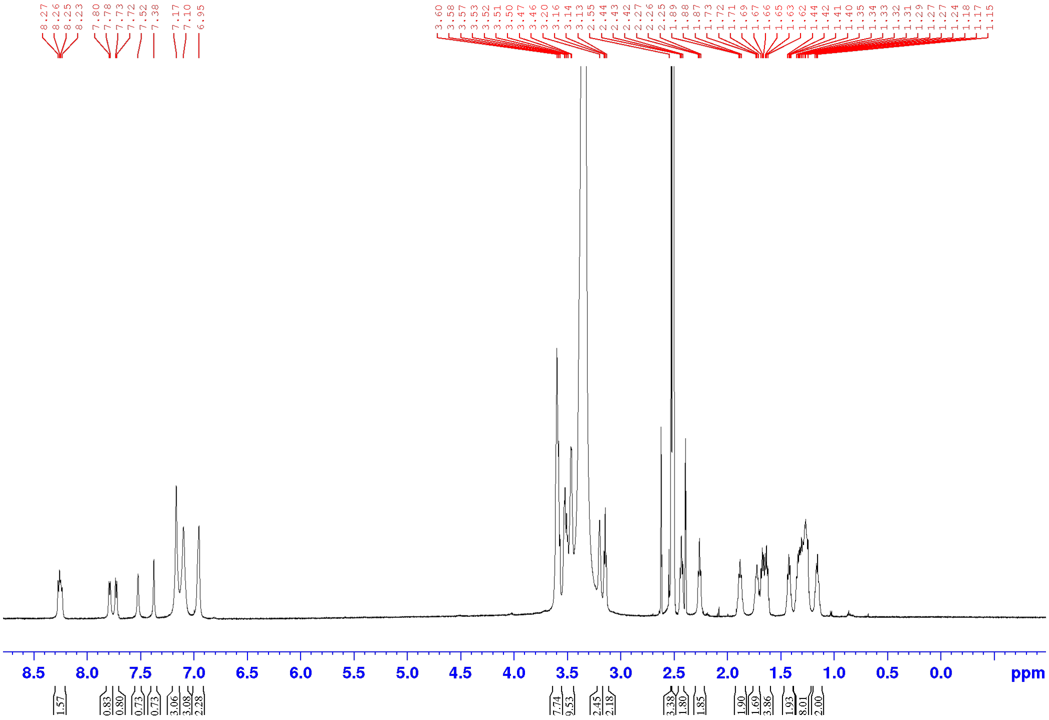

#### 5.2 3-(3-(7-(Bis(methyl-d3)amino)-3-(bis(methyl-d3)iminio)-5,5-dimethyl-3,5-dihydrodibenzo[*b,e*]silin-10-yl)-4-carboxy-*N*-(2-(2-((6-chlorohexyl)oxy)ethoxy)ethyl)benzamido)propane-1-sulfonate (SiR-d12-HTL)

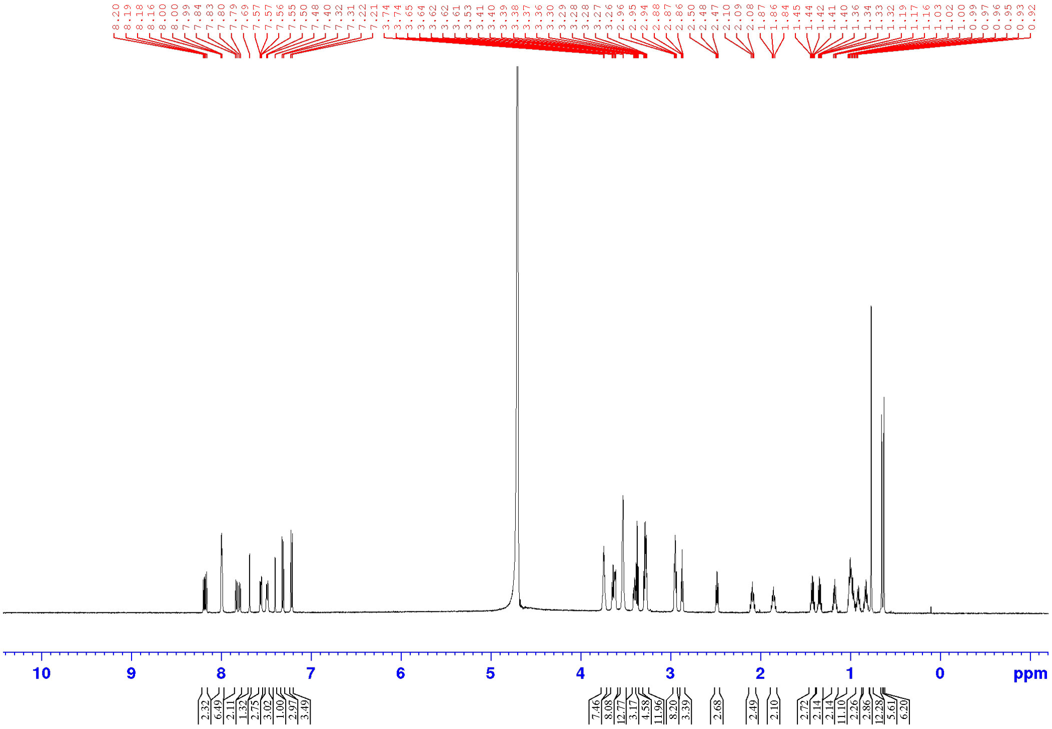

#### 6 Computational modelling

The ligand structures of TMR-SHTL and TMR-HTL were geometry optimised in isolation using the MMFF94 force field with a conjugated gradient algorithm.[5] The protein receptor structure, as well as the native covalent ligand structure, was obtained from the RCSB protein database (PDB-ID 6Y7A).[6]

After receptor preparation, including removal of heteroatoms and protonation adjustment for physiological conditions, the ligand structures were fitted covalently to the receptor using a custom script based on the two-point attractor method.[7] The active residue (ASP106) was set to have a flexible side-chain and covalent ligand-receptor docking was performed using the AutoDock 4 engine.[8] The docking system was set up using a 32 Å × 32 Å × 32 Å centroid around the active residue of the receptor protein and by calculating 3 binding modes with exhaustiveness of 32. The computation was performed 10 times for analysis of descriptive statistics (**Supplementary Figure S4**).

The structural deviation from the reference ligand (TMR-HTL) was calculated as the RMSD, as well as the atom-resolved positional deviation, as depicted in **Supplementary Figure S3**.

Calculation of the significance level of binding affinity deviation of the best poses was performed with respect to the reference ligand (TMR-HTL) using an exact Mann-Whitney test at a two-tailored confidence level of 95 %.[9]

**Mann-Whitney:** $U = 32$, $p = 0.1903$.

### 7 Supplementary Figures

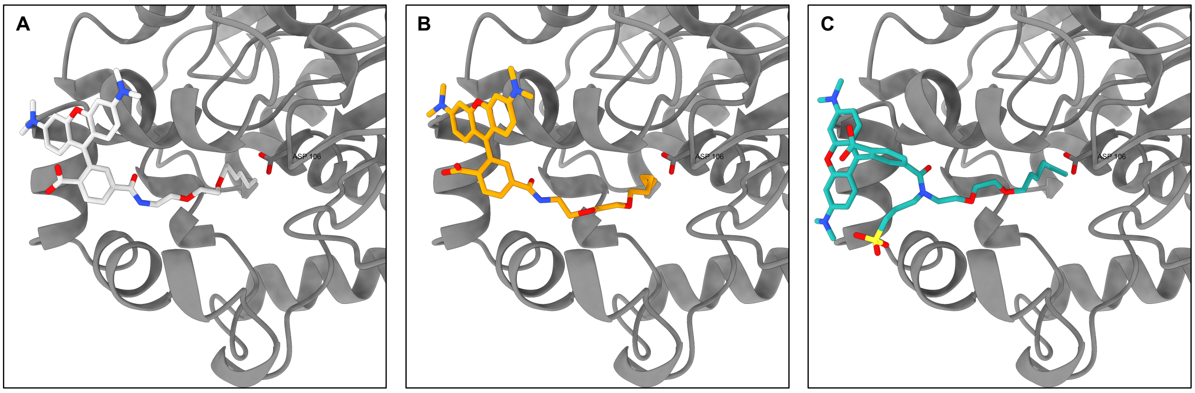

**Supplementary Figure S1: Modelled Covalent Docking Poses.** The computed best-scoring dock poses of both ligands are compared to the crystal structure reference ligand. **A** Reference structure HTP:TMR. **B** Modelled HTP:TMR. **C** Modelled HTP:S-TMR.

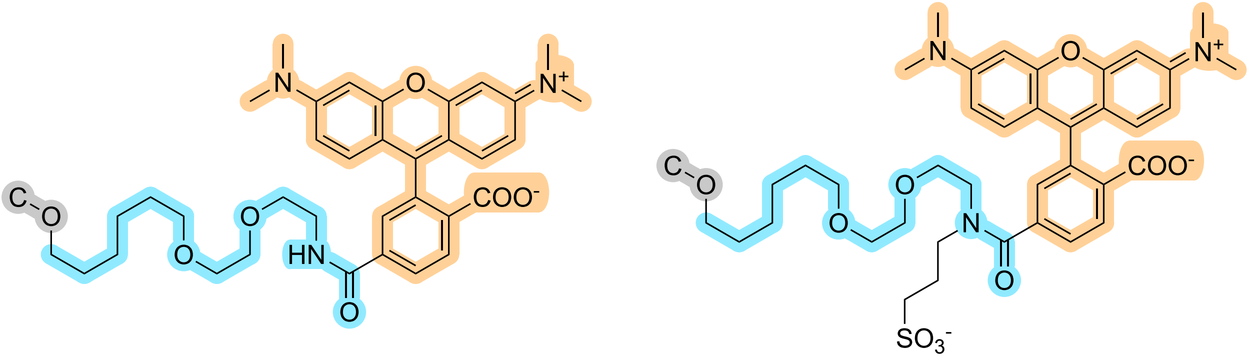

**TMR-HTL** **TMR-SHTL**

**Supplementary Figure S2: Investigated Ligand Structures.** The highlighted regions in both structures were used to compute the RMSD values for all modelled systems (grey: two-point covalent attractor, blue: HaloTag-linker, orange: fluorophore).

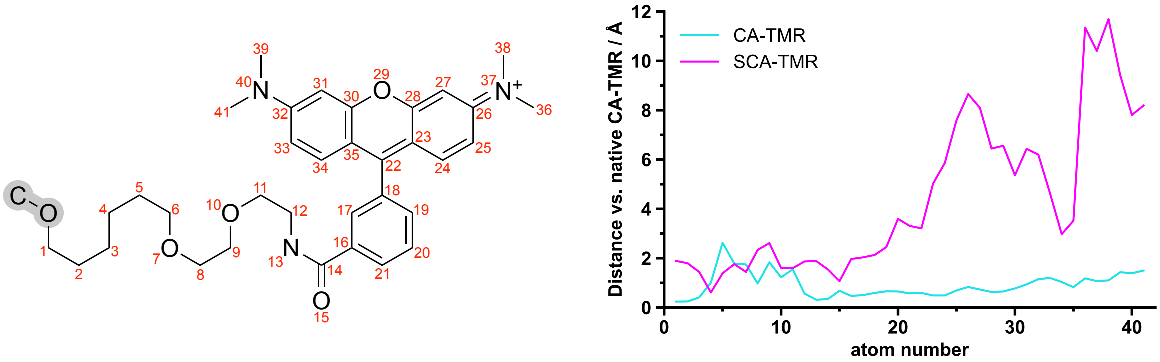

**Supplementary Figure S3: Structural Deviation from the Reference Ligand.** The structural motif present in both TMR-HTL and TMR-SHTL is depicted, as well as the two-point attractor (grey). Numbering of the atoms is coherent with the distance plot, showing the structural flexibility of the unordered HaloTag-linker in all structures, as well as the close structural agreement between the reference ligand and the modelled TMR-

HTL, whereas the modelled TMR-SHTL deviates more significantly from the reference structure.

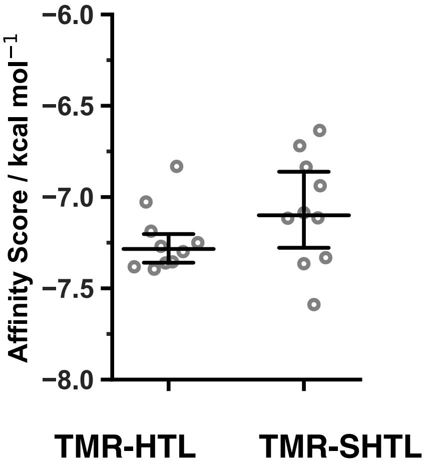

**Supplementary Figure S4: Binding Affinity Scores.** Comparison of 10-fold computation results for HTP:HTL-TMR and HTP:SHTL-TMR. The difference is not significant ($p = 0.1903$), displayed are all scores as well as median, 25 % and 75 % quartile bars.

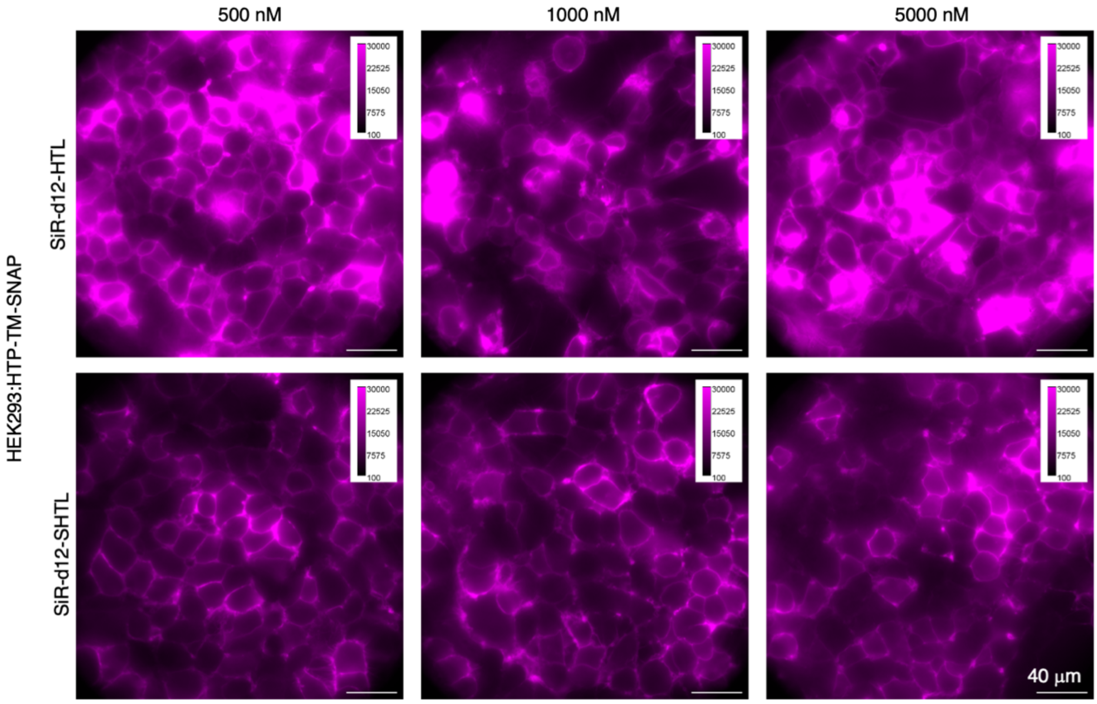

**Supplementary Figure S5: Titration experiment using SiR-d12-HTL and SiR-d12-SHTL on live HTP-TM-SNAP transfected HEK293 cells.**

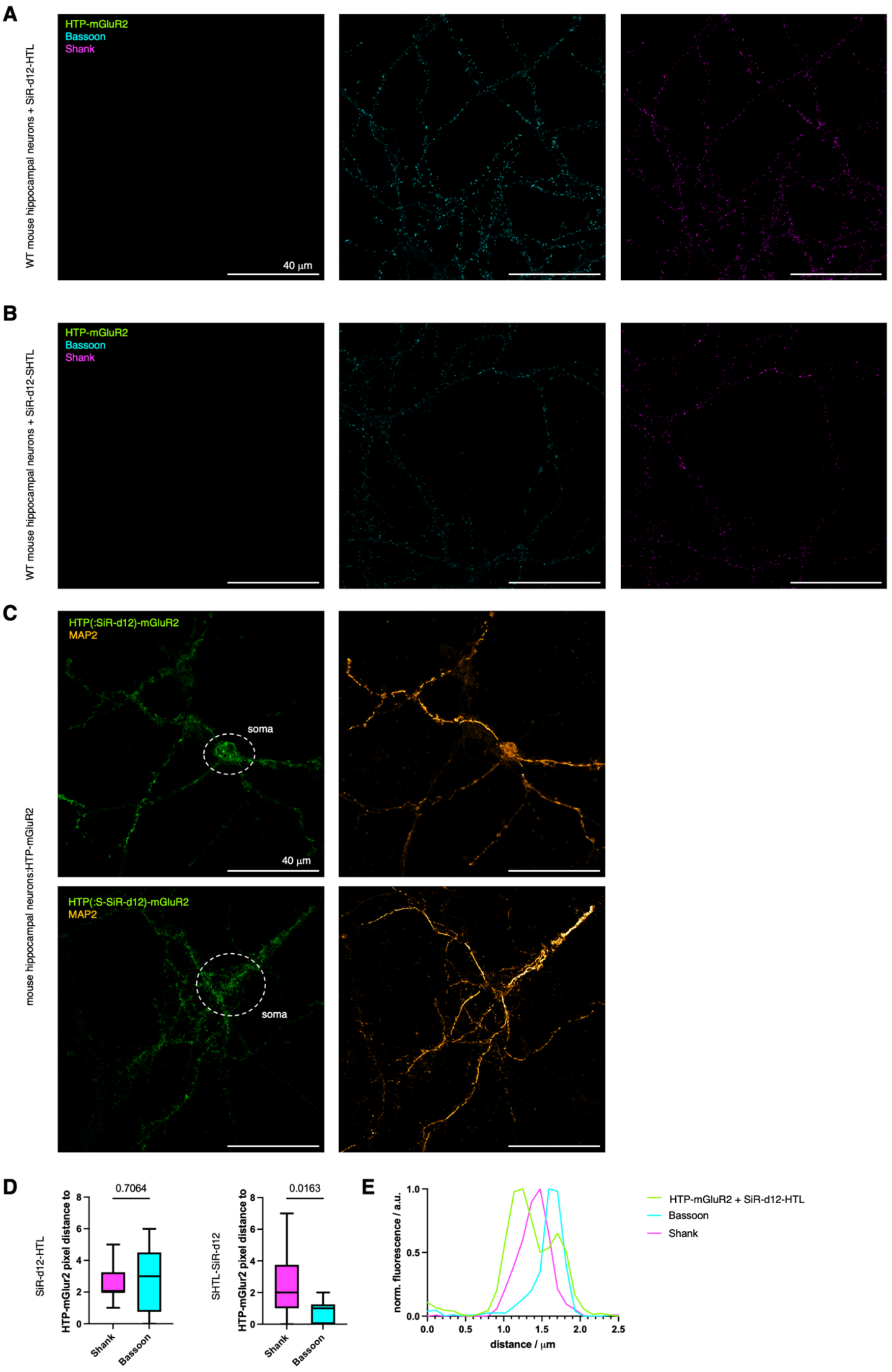

**Supplementary Figure S6:** Non-transduced neurons stained with SiR-d12-HTL **(A)** and SiR-d12-SHTL **(B). C)** MAP2 and HTP-mGluR2 single channel images from Figure 3D. **D)** Line scans of synapses using SiR-d12-HTL, similar to Figure 3K (on right hand side for easy comparison, using SiR-d12-SHTL. **E)** Representative line scan profile from D with higher intracellular mGluR2 localization.

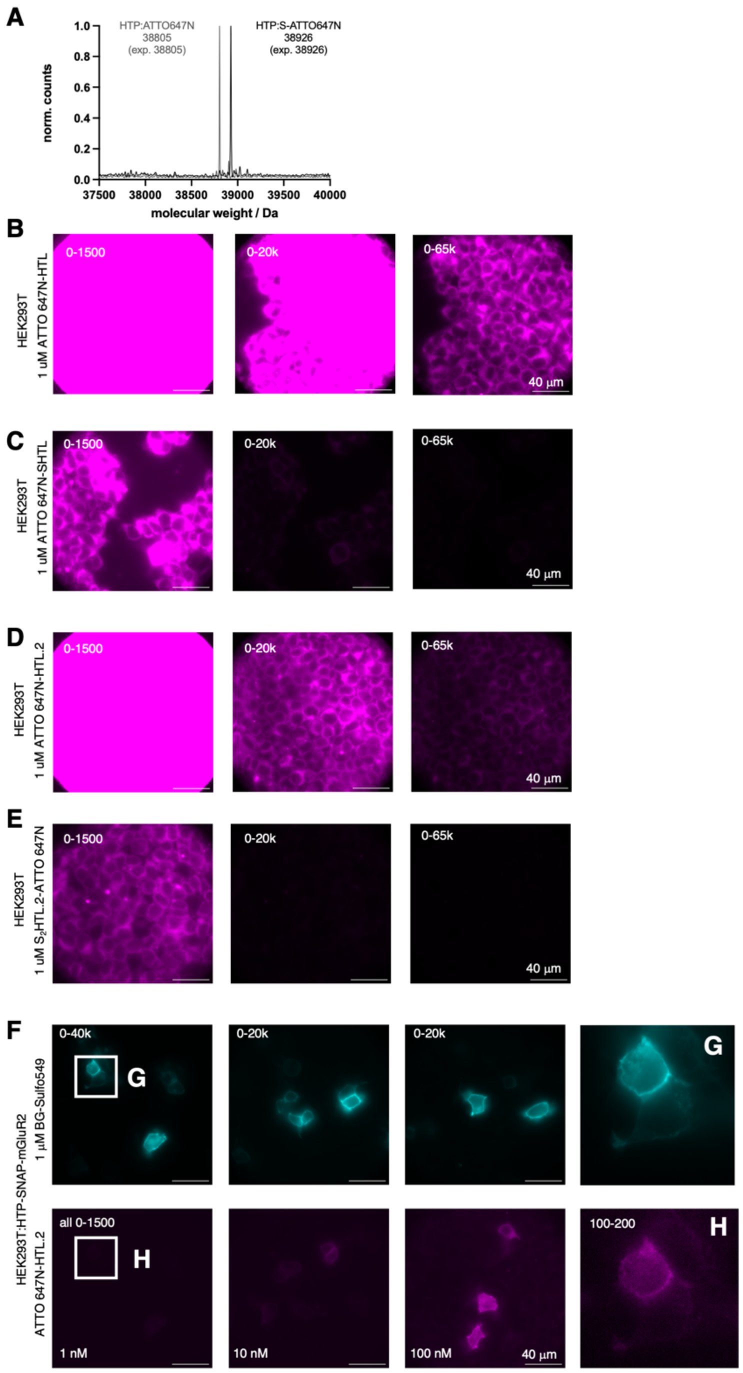

**Supplementary Figure S7: A)** Full protein mass spectrometry shows HTP-labelling with ATTO 647N-HTL and ATTO 647N-$S_2$HTL.2. **B-E)** HEK293 cells treated with 1 µM ATTO 647N-HTL (B), SHTL (C), HTL.2 (D) and $S_2$HTL.2 (E) with different brightness and contrast settings for comparison. **F)** SNAP-HTP-mGluR2 transfected HEK293 cells treated with BG-Sulfo549 (1 µM) and titrated with ATTO 647N-HTL. **G)** Zoom in of cells treated with BG-Sulfo549. **H)** As for (G) but with ATTO 647N-$S_2$HTL.2.

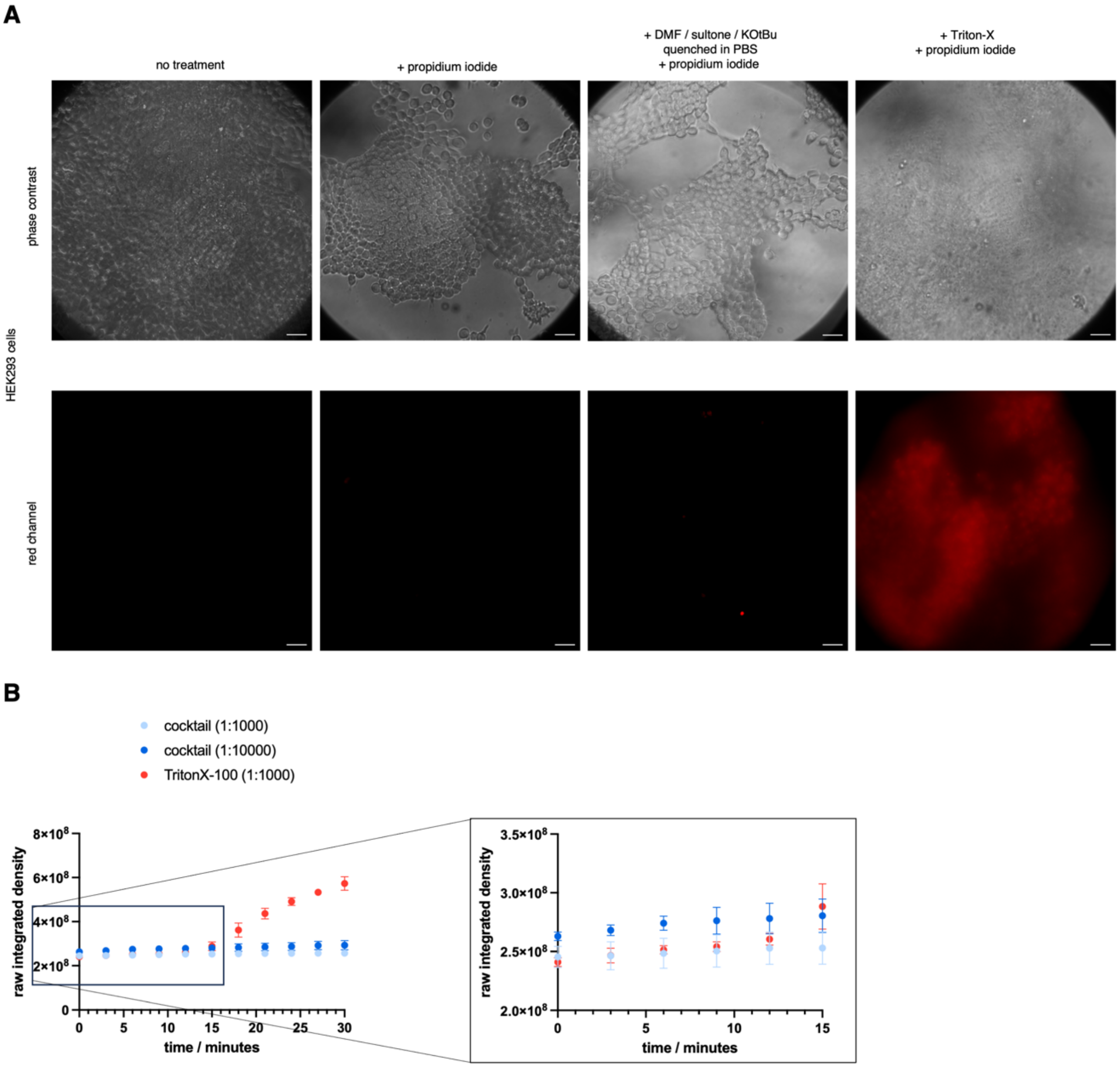

**Supplementary Figure S8: PI assay using the in situ cocktail with popidium iodide. A)** Widefield imaging of HEK293 cells after 5 minutes incubated with propidium iodide (1:250 of a 9 wt% solution), and additional cocktail or TritonX-100 as positive control. Scale bar = 40 micrometer. **B)** Integrated density of propidium iodide (1:1000 of a 9 wt% solution) over time shows cell death after a15 minutes, while different dilutions of the labelling cocktail does not lead to increased signal intensity. n = 3. Mean±SD.

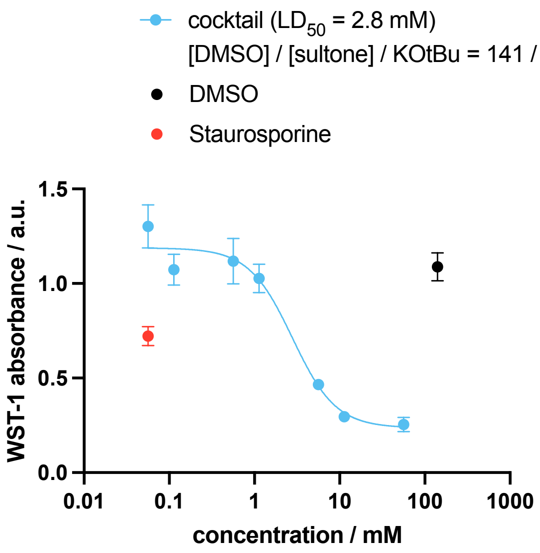

**Supplementary Figure S9: WST-1 assay.** HEK293T cells were seeded (40,000 cells/well) in a clear 96 well-plate and allowed to grow for 1 days in 100 uL DMEM supplemented with 10% FBS at 37 °C and 5% $CO_2$. Solutions were prepared in full medium, cells were aspirated before addition of the solutions. Incubation occurred over night. WST-1 (#MK400, Takara Bio) was added on top according to the manufacturer's instructions. Incubation was performed for another 2 hours before absorbance was read on a TECAN INFINITE M PLEX plate reader ($\lambda_{Abs}$ = 440 nm) and corrected by subtraction ($\lambda_{Abs\ correct}$ = 660 nm). Plotting was performed in GraphPad Prism 8. n = 3-6. Mean±SD.

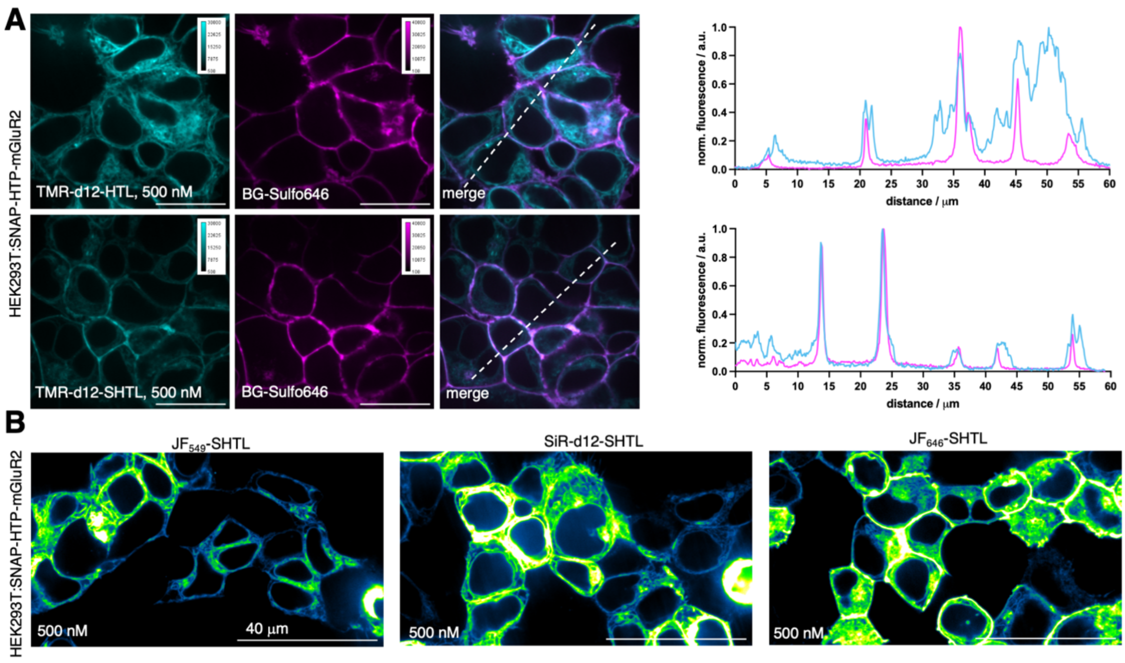

**Supplementary Figure S10: Staining as for Figure 6, but with 1:1000 fold dilution of the reaction mixture shows less clear surface staining.**

#### 8 Supplementary Tables

**Supplementary Table 1: RMSD Calculations.** The computational results are compared to the native ligand present in the experimental crystal structure of the receptor protein.

| | TMR-HTL | TMR-SHTL |
|---|---|---|
| **RMSD vs. native TMR-HTL** | 1.047 Å | 5.368 Å |

**Supplementary Table 2: Covalent Docking Scores.** The investigated HTP:TMR and HTP:S-TMR systems were modelled independently (randomised seed) 10 times. Depicted are some descriptive parameters for the achieved best-fit docking affinity scores for both systems (all in kcal/mol).

| | TMR-HTL | TMR-SHTL |
|---|---|---|
| **Number of values** | 10 | 10 |
| **Minimum (best score)** | -7.395 | -7.589 |
| **Maximum** | -6.832 | -6.635 |
| **Mean** | -7.236 | -7.073 |
| **Std. Deviation (corr.)** | 0.179 | 0.301 |

#### 9 One-step reaction protocol

Please cite Roßmann et al., 2024.

We recommend to work in a fume hood.

- Weigh in 1.0 mg KO*t*Bu (Aldrich: #659878) into an Eppendorf tube.
- Add 104 uL of DMSO (we recommend single use ampules: e.g. Carl Roth: #AE56.3, 10 x 0.75 mL) to make a 100 mM stock solution.

- Add 1,3-propane sultone into an Eppendorf tube and warm on a shaker to 35 °C to obtain a liquid (mp = 32 °C)
- Add 4 uL of DMSO solution to 5 nmol of the dye-HTL conjugate (e.g. Promega: #HT1020 for $JF_{549}$-HTL and #HT1060 for $JF_{646}$-HTL – in this case the 5 x 1 nmol aliquots can be pooled), briefly vortex to ensure full dissolution and spin down.
- Add 1 uL of liquid sultone into the DMSO solution and pipette up and down to ensure full mixing.
- Allow incubation for 5 minutes at room temperature.

- Add 5 uL PBS to the reaction mixture and briefly vortex and spin down.
- The reaction is completed and quenched. Dilute 1:10,000 in medium of choice, aspirate cells medium and replace with labelling solution for 30 minutes at 37 °C. Wash cells once and proceed with protocol of choice (e.g. fixation, imaging).

<u>Concentrations during the steps:</u>

[DMSO] = 14.1 M

[1,3-propane sultone] = 10.8 M

[KO*t*Bu] in DMSO = 100 mM

Adding sultone to DMSO:

[DMSO] = 11.3 M

[1,3-propane sultone] = 2.16 M

[KO*t*Bu] = 80 mM

Quench with PBS, hydrolyzing sultone to sulfonic acid:

[DMSO] = 5.64 M

[3-hydroxypropanesulfonic acid] = 1.08 M

[KO*t*Bu] = 40 mM

Dilute 1:10000:

[DMSO] = 564 μM

[3-hydroxypropanesulfonic acid] = 108 μM

[KO*t*Bu] = 4 μM
